# Predictive framework of macrophage activation

**DOI:** 10.1101/2021.08.02.454825

**Authors:** David E. Sanin, Yan Ge, Emilija Marinkovic, Agnieszka M. Kabat, Angela Castoldi, George Caputa, Katarzyna M. Grzes, Jonathan D. Curtis, Sebastian Willenborg, Stefanie Dichtl, Susanne Reinhardt, Andreas Dahl, Erika L. Pearce, Sabine A. Eming, Alexander Gerbaulet, Axel Roers, Peter J. Murray, Edward J. Pearce

## Abstract

Macrophages populate every organ during homeostasis and disease, displaying features of tissue imprinting and heterogeneous activation. The disjointed picture of macrophage biology that emerged from these observations are a barrier for integration across models or with *in vitro* macrophage activation paradigms. We set out to contextualize macrophage heterogeneity across mouse tissues and inflammatory conditions, specifically aiming to define a common framework of macrophage activation. We built a predictive model with which we mapped the activation of macrophages across 12 tissues and 25 biological conditions, finding a striking commonality and finite number of transcriptional profiles, which we modelled as defined stages along four conserved activation paths. We verified this model with adoptive cell transfer experiments and identified transient RELMɑ expression as a feature of macrophage tissue engraftment. We propose that this integrative approach of macrophage classification allows the establishment of a common predictive framework of macrophage activation in inflammation and homeostasis.

**One Sentence Summary:** We propose an integrative approach of macrophage classification that allows the establishment of a common framework of macrophage activation in inflammation and homeostasis.

## Introduction

Macrophages can be found in every organ and displaying a unique transcriptional profile in each setting (*1, 2*). This profound specialization to match their tissue of residence is a necessary aspect of the function of these cells during homeostasis (*3*). The engraftment of macrophages in most tissues occurs early during embryonic development (*4, 5*). In adults, circulating monocytes contribute to the replenishment of these tissue-resident macrophage pools at different rates, or not at all, depending on the organ in question (*5–9*). The extent of the contribution of monocytes to tissue macrophage populations during adulthood is an area of debate and findings from newly developed fate-mapping tools require ongoing revision of macrophage ontogeny models (*4, 6, 10*). These observations have only recently been expanded to humans (*11*). However, the relatively simplistic view held for decades after the introduction of the mononuclear phagocyte system (*12*), in which all macrophages differentiate from bone-marrow derived monocytes, has been abandoned.

As focus has shifted to the origin of macrophages and the impact of tissue imprinting (*1*), the engagement of recruited versus resident macrophages during the immune response has received greater scrutiny. However, these efforts have been hampered by the limitations inherent to phenotyping techniques that rely on bulk population averaging (e.g. RNA sequencing), few simultaneous measurements (e.g. flow cytometry) and poorly characterised macrophage subset markers. This is especially challenging as incoming monocytes are able, with time, to adopt nearly indistinguishable transcriptional profiles to resident macrophages in the tissue that they are entering (*13*). Despite these limitations, some have taken the sum of these studies to suggest that macrophages in different tissues should be regarded as entirely different cells (*3*), or that paradigms of macrophage M1 (classical)/M2 (alternative) activation should be abandoned (*14–16*). This later view in particular is supported by the extensive plasticity that macrophages display when stimulated with cocktails of cytokines, pattern recognition receptor ligands and other immunomodulatory molecules (*17, 18*). Thus, the emerging picture of macrophage activation suggests a flexible spectrum of different activation states, with tissue and context-specific parameters viewed as dominant predictors of macrophage function.

This complex landscape of macrophage phenotype has been further thrown into relief by the emergence of single cell RNA sequencing (scRNA-seq). This technology overcomes the limitations of bulk population averaging and does not rely on previously defined surface markers for macrophage subset sorting. As more studies employing this technique are published, the observed heterogeneity in macrophage activation states has further increased, with new subsets or phenotypes frequently identified (*19–28*). Consequently, the field of macrophage biology currently lacks a common reference framework to describe the state of activation of macrophages in tissues.

In light of this rapidly evolving situation, we wondered whether the construction of such a common framework would be possible. We reasoned that a unifying model could be built by comparing macrophage activation profiles across tissues under multiple inflammatory conditions. We expected that either we would succeed in finding common activation features or that tissue-specific transcriptional programs would dominate the data. With this in mind, we built a predictive model with which we mapped the activation of macrophages across 12 mouse tissues and 25 biological conditions, finding a strikingly common and finite number of transcriptional profiles which we modelled as stages along 4 conserved activation paths. These activation stages placed cells with varying frequencies along a “phagocytic” regulatory path, an “inflammatory” cytokine producing path, an “oxidative stress” apoptotic path or a “remodelling” extracellular-matrix (ECM) deposition path. We verified our model with adoptive cell transfer experiments, noting that incoming monocytes displayed a remarkable plasticity to rapidly adopt all the transcriptional signatures we detected. Moreover, we identified transient RELMɑ expression as a feature of macrophage tissue engraftment and propose that historical RELMɑ expression may serve to identify monocyte contribution to tissue resident macrophage populations. Lastly, we posit that this integrative approach of macrophage classification allows the establishment of a common predictive framework of macrophage activation that may serve to contextualize these cells in future studies and for this reason we provide a list of surface markers that may be used to identify these cells. For this purpose, we built a web interface where interested researchers may explore our findings interactively (https://www.macrophage-framework.jhmi.edu).

## Results

### Macrophages in inflammatory conditions co-exist in diverse functional states

scRNA-seq has highlighted extensive heterogeneity in macrophage populations across tissues and conditions (*5, 11, 22, 24, 27, 29*), and the picture of macrophage biology that has emerged from these studies can be difficult to integrate with the paradigms of macrophage activation that have developed from *in vitro* studies. For this reason, we set out to contextualize macrophage heterogeneity across tissues in diverse inflammatory conditions, specifically aiming to define common aspects of macrophage activation during infection and inflammation (Figure 1A). For this purpose we built a reference dataset (Figure 1A-B) based on 2 inflammatory conditions historically seen as representing either a classical inflammatory response during bacterial infection using *Listeria monocytogenes* (*L. mono*), or a type-2 immune response to *Heligmosomoides polygyrus* (*H. poly*) larvae. Given our goal of encapsulating most macrophage activation states, the completeness of our reference dataset was critical. We reasoned that the two settings we chose, which both induce multi-cellular and systemic responses, would provide a sufficiently broad spectrum of macrophage activation to begin our study.

**Fig. 1.**
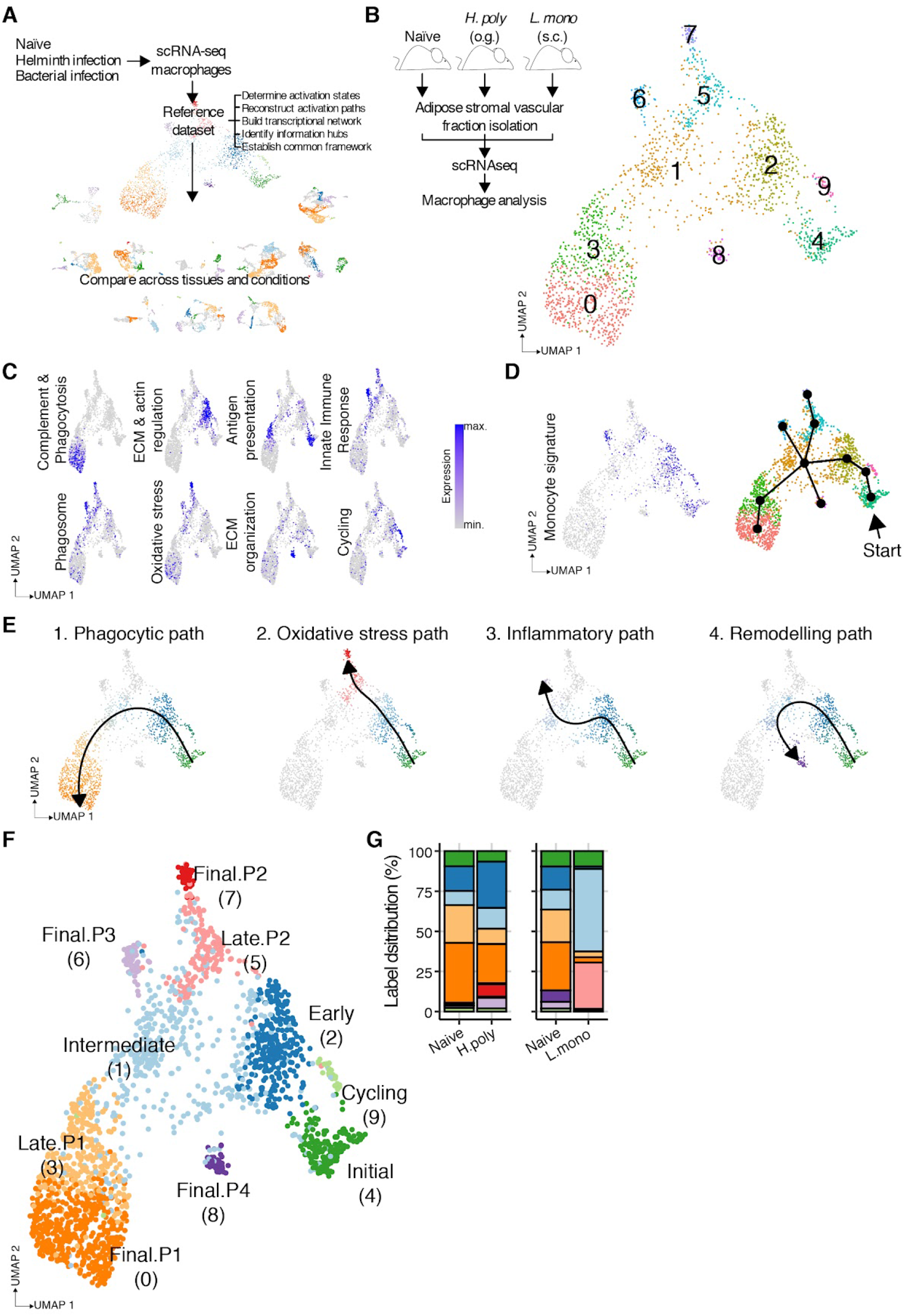
Macrophage activation in inflamed tissues follows predefined paths. (A) Schematic depiction of reference dataset construction, outlining overall goals of strategy. (B) scRNA-seq analysis of macrophages (cells = 2000) from the stromal vascular fraction (SVF) of adipose tissue from naïve, *H. poly* or *L. mono* infected animals (n = 1-8 per group) shown as a UMAP, highlighting identified clusters. (C) Relative levels (low - gray; high - blue) of gene set scores associated with identified clusters. (D) Relative levels (low - gray; high - blue) of gene set scores associated with Monocyte signature (left) and predicted lineage breaking points (right). (E) Lineage and pseudotime calculation showing activation trajectories. Cells assigned to identified paths are colored to match stage labels. Non-participating cells are shown in gray. (F) UMAP labelled according to path progression indicating shared (Initial > Early > Intermediate) and path specific (Phagocytic: Late.P1 > Final.P1; Oxidative stress: Late.P2 > Final.P2; Inflammatory: Final.P3; Remodelling: Final.P4) macrophage activation stages. Cluster number indicated in brackets. (G) Activation stage distribution shown as a percentage of total cells per biological condition.

Initially, we sequenced all stromal vascular fraction cells from mesenteric fat (Figure S1A), adjacent to the site of *H. poly* infection, and popliteal fat (Figure S1B), which is directly invaded by *L. mono* following footpad injection. We then evaluated the distribution of macrophage gene expression markers in these datasets (Figure S1A-B) and extracted, integrated and re-clustered identified macrophages. To ensure a balanced representation of each condition, only 500 macrophages were taken from each dataset, plus 500 macrophages from matched naive controls. Gene expression was distinct within each identified cluster (Figure S1C & Supplemental Table 1) and could be associated with specific biological processes via pathway enrichment analysis (Figure 1C). To better visualize gene expression programs we calculated gene set scores within each cell for groups of genes mapping to diverse functions (Supplemental Table 2). These gene set scores were specific to different identified clusters, thus underscoring the functional diversity of sequenced macrophages in the conditions studied.

We observed that clusters 0 and 3 were enriched for genes associated with macrophage alternative activation (e.g. *Cd36, Clec10a, Mrc1*), antigen presentation (e.g. *H2-Aa, H2-Eb1, H2-Ab1*) and the complement cascade (e.g. *C1qc, C1qb*) (Figure 1C, S1C & Supplemental Table 1). Cluster 2 was enriched for genes involved in extracellular matrix (ECM) receptor-interactions (e.g. *Cd44, Sdc1, Fn1*) and cytoskeleton regulation (e.g. *Pfn1, Actg1, Tmsb4x*). Cluster 4 displayed high expression of genes participating in antigen presentation (e.g. *H2-Oa, H2-DMb2, Cd74*). Clusters 6 and 7, and to a lesser extent 5, were enriched for genes associated with the phagosome (e.g. *Fcgr1, Ncf4, Fcgr3*) and oxidative stress (e.g. *Prdx5, Txn1, Gsr*), with cluster 6 in particular enriched for innate immune response genes (e.g. *Ifitm3, Fcgr1, Isg20*). ECM organization genes (e.g. *Col1a1, Col3a1, Ddr2*) were highest in Cluster 8, while Cluster 9 displayed high expression of cell cycle associated genes (e.g. *Cks1b, H2afx, Cks2*). Interestingly, all but one cluster could be assigned a functional specialization in the manner described above. Cluster 1, which also occupied the center of the UMAP, had no distinctly regulated genes (Figure S1C & Supplemental Table 1) and therefore no pathway assignment (Figure 1C). Thus, our analysis shows that macrophages within a tissue simultaneously specialize into multiple functional stages, echoing findings in other studies where this diversity has been reported (*22, 24, 27*).

### Macrophages in inflammatory conditions are arranged along activation paths

In our data, cluster 1 could not be associated with a distinct function as it displayed no up-regulated genes using the thresholds we established (average log fold change > 1, percent cells expressing gene > 0.4 and an adjusted p value < 0.01). Despite these high stringency filters, we reasoned that cluster 1 could in fact represent an intermediate state of activation, suggesting that rather than different populations of macrophages, our transcriptional analysis captured activation paths being followed by infiltrating macrophages. To address this hypothesis we analysed our data with Slingshot (*30*) to calculate first lineage breaking points (Figure 1D) and later lineage curves associated with pseudotime (Figure 1E), generating a model of macrophage activation in the tissue. This analysis required the selection of a starting point for the curves. To establish this starting point, we calculated a gene set score associated with monocytes (Figure 1D & Supplemental Table 2) based on their transcriptional profile (*1*). Given the reported increase in Major histocompatibility complex class II (MHC-II) in infiltrating monocytes transitioning to macrophages (*31*) and the increased monocyte signature we observed, we reasoned that cluster 4 (Figure 1B-D, black arrow) was a suitable starting point for our activation model.

Our analysis identified 4 activation paths that we labelled as “Phagocytic”, “Oxidative stress”, “Inflammatory” and “Remodelling” according to the enriched pathway at the end point clusters of each lineage (Figure 1E). Our analysis also revealed that at least three clusters (4, 2, 1) represented common early stages of macrophage activation. For clarity, we named these early stages of macrophage activation according to their relative position in the pseudotime progression as “Initial” (cluster 4), “Early” (cluster 2) and “Intermediate” (cluster 1) stages. The remaining clusters were renamed as either “Late” or “Final”, with the activation path appended at the end of the stage (Figure 1F), so that for instance “Late.P1” corresponded to the cluster in activation path 1 (P1) that is between the “Intermediate” and “Final” stage.

Next we investigated how these activation stages were distributed across the biological conditions in the data (Figure 1G). We observed that the distribution of naive cells into different functional stages in different fat deposits was comparable, and diverse, perhaps indicating an active process of macrophage activation as monocytes replenish these sites in steady state (Figure 1G). We also noted that both helminth and bacterial infections altered the proportions of several of these stages, such that *L. mono* infection favored the “oxidative stress” path, while *H. poly* infection favored the “phagocytic” path when compared to each other (Figure 1G). Notably, all stages were present in each condition, underscoring the difficulty of relying on bulk phenotyping techniques to capture the overall picture of macrophage activation *ex vivo*. Moreover, the changes in diversity induced by infection were notably different between the inflammatory conditions studied. While *H. poly* infection resulted in increases in numbers of cells within less-well represented stages, *L. mono* infection resulted in 2 stages becoming dominant. These differences could be explained both by the specific immune responses tailored to the pathogens involved, but also by the time at which samples were collected (Supplementary Table 3; day 1 p.i, for the *L. mono* dataset, day 14 p.i. for the *H. poly* dataset).

In summary, our model predicts that macrophages in an inflamed tissue are progressing through several distinct activation stages with unique transcriptional profiles. Moreover, the balance of this progression is influenced by the type of immune response that dominates the microenvironment but perhaps also the timing of this response. Finally, our model suggests that although a tissue can become dominated by relatively few activation stages, there are still macrophages present that have committed to other paths.

### Macrophage gene expression is regulated along activation paths

Our initial analysis of macrophage activation relied on comparing gene expression in each cluster to all remaining cells in the dataset (Figure S1C). We complemented this analysis with a different approach, where gene expression was modelled as a function of pseudotime (Figure 2A). We reasoned that as macrophages progress along each of the activation paths we defined (Figure 1E-F), gene expression would be regulated to allow these cells to become fully functional. As we thought it would be unlikely that the relationship between gene expression and pseudotime would be linear, we instead fitted a general additive model (GAM), using non-parametric locally estimated scatterplot smoothing (loess), explaining the expression of a gene as a function of the relative position of a cell along an activation path (Figure 2A). We included in this analysis only the top 2000 most variable genes in the cells of each path, and ranked the resulting models based on the p value of the association of pseudotime and gene expression (Figure S2A). We found that collectively the expression of 828 genes (p value < 1x10^9^) could be modelled in this way and we show the top most significant association for each pathway (Figure 2B). Moreover, we observed that the regulation of typical macrophage activation markers followed expected behaviours in our modelling approach and corresponded to defined activation paths (Figure 2C & S2B). For instance, only cells in the “phagocytic” path exhibited a steep and continuous increase in alternative activation markers (*Il4ra*, *Mrc1*, *Clec10a*) and the mitochondrial metabolism gene *mt-Co1* as a function of pseudotime (Figure 2C). Conversely, expression of inflammatory genes (*Il6*, *Il1b*) was only retained at high levels in the “inflammatory” and “remodelling” paths (Figure 2C).

**Fig. 2.**
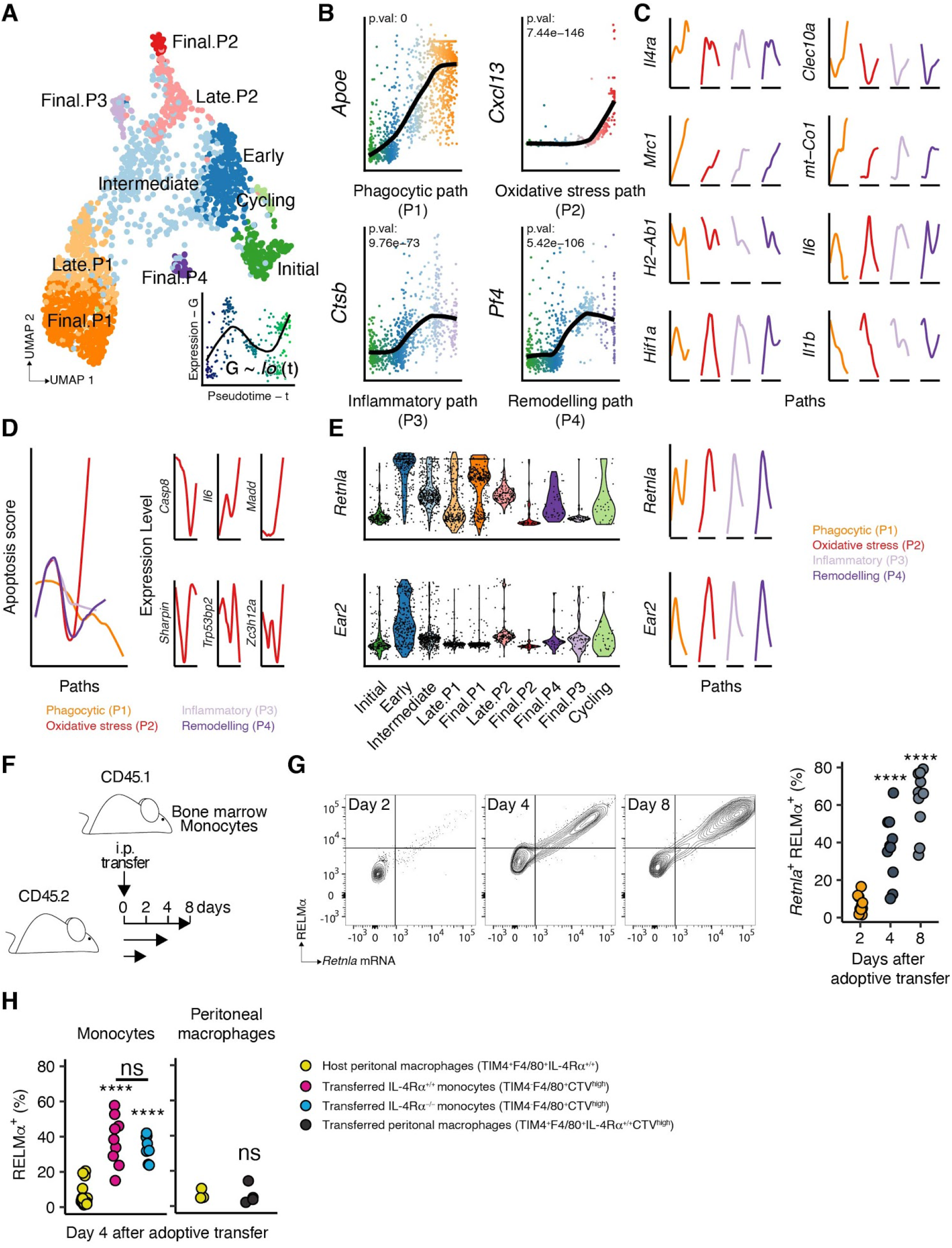
Macrophage gene expression can be modelled as a function of activation, revealing a transitory stage of RELMɑ expressing cells. (A) Macrophage activation stage UMAP, showing an example of a fitted general additive model (GAM) for gene expression as a function of pseudotime. (B) Top most significant GAM fits for genes associated with identified paths, showing single cells gene expression (dots - color matching activation stage) as a function of pseudotime (x axis) for each activation path, with fitted models (black lines) and associated adjusted p values also shown. (C) Fitted GAM models (colored lines matching activation paths) of gene expression of macrophage activation markers (y axis - fixed across all paths for each gene) as a function of pseudotime (x axis - specific to each path). (D) Left - Fitted GAM models (colored lines matching activation paths) of Apoptosis score (y axis) as a function of pseudotime (x axis). Right - Fitted GAM model of expression level (y axis) of a subset of genes used in score calculation as a function of pseudotime (x axis) shown for “oxidative stress” path. (E) Relative expression of RELMɑ coding gene *Retnla* and *Ear2* shown as violin plots for each activation stage (left) or as a fitted GAM models specifically for each activation path (right). (F) Schematic view of experimental set up for adoptive bone marrow monocyte transfer from donor (CD45.1^+^) mice into the peritoneal cavity of naïve recipients (CD45.2^+^). (G) Representative density flow cytometry plots (left) and quantification of percentage positive cells (right) expressing RELMɑ protein and mRNA (*Retnla*) in adoptively transferred bone marrow monocytes recovered from the peritoneal cavity of recipient mice at indicated times. Individual dots represent biological replicates from combined experiments (n = 10 across 2 repeats). Significant differences at each stage compared to day 2 are indicated (****: p value < 0.0001) based on single factor anova analysis followed by Tukey Honest Significant Differences test. (H) Quantification of percentage positive macrophages expressing RELMɑ 4 days post adoptive cell transfer within host large peritoneal macrophages, adoptively transferred IL-4Rɑ sufficient (IL-4Rɑ^+/+^) or deficient (IL-4Rɑ^-/-^) monocytes (left), or in adoptively transferred large peritoneal macrophages (right). Individual dots represent biological replicates (n = 4-18). Significant differences at each stage compared to host macrophages are indicated (**: p value < 0.01; *: p value < 0.05; ns: p value > 0.05) or between transferred cells (ns: p value > 0.05) based on single factor anova analysis followed by Tukey Honest Significant Differences test.

We next reasoned that not only individual genes but also gene set scores could be modelled in this manner. Consequently, we calculated the aggregate expression of genes associated with apoptosis (Supplemental Table 2) and visualized the regulation of this gene expression program across activation paths (Figure 2D). Interestingly, cells at the end of the “oxidative stress” path displayed the highest levels for the apoptosis score (Figure 2D, red lines), while cells at the end of the “phagocytic” path had the lowest (Figure 2D, orange line). In addition to aligning with reports demonstrating that macrophage activation can result in cell death (*32–34*), these results could indicate that commitment to an activation path might be unidirectional: in other words, macrophages would not transition from one path to another, although this would need to be demonstrated experimentally. Finally, these results indicate that cells committed to the “phagocytic” path, might be long-lived and go on to replace tissue resident cells. Collectively, our results show that the proposed activation model broadly agrees with published expectations of macrophage gene expression regulation and offers the possibility to uncover new aspects of macrophage biology.

### Macrophages transition through a RELMɑ expressing activation stage

Defining trajectories based on scRNA-seq data greatly depends on the dimensional projection upon which the analysis is based. Consequently, we sought to validate the activation model by exploiting our gene expression analysis approach to extract markers of an intermediate stage defined in our results. Our initial exploration of the data revealed that the RELMɑ encoding gene *Retnla* was found both in cells committed to the “phagocytic” path (P1) (Figure 2E - left), which expressed several other markers of alternative macrophage activation (Figure 2C and Supplemental Table 1), and also in the “Early” activation stage shared by all paths (Figure 2E - left). Exploring the relationship between *Retnla* expression and pseudotime, we observed that our model predicted a wave of expression early during activation (Figure 2E - right). We observed also that this was not the case for all genes expressed in this stage and show that *Ear2* expression, like *Retnla*, peaks at this early stage, but then steadily drops (Figure 2E).

We reasoned that if our model was correct, then monocytes would start expressing *Retnla* shortly after entering a tissue; this would occur regardless of the inflammatory state of that tissue. To test this, we isolated bone marrow monocytes from CD45.1^+^ mice (Figure S2C), and adoptively transferred them into the peritoneal cavity of CD45.2^+^ naive hosts (Figure 2F). We then evaluated the abundance of *Retnla* mRNA and RELMɑ protein in CD45.1^+^ macrophages recovered on days 2, 4 and 8 days after adoptive transfer (Figure 2G and S2D). Confirming the model’s predictions, we observed that macrophages that had differentiated from transferred monocytes began to express RELMɑ 4 days post-adoptive transfer and that expression continued to increase until ∼80% of the cells were positive for this molecule (Figure 2G). RELMɑ induction occurred in the absence of IL-4Rɑ stimulation, as we did not detect changes in RELMɑ expression in the resident cells in the peritoneal cavity and moreover IL-4Rɑ^-/-^ monocytes displayed similar RELMɑ expression compared to IL-4Rɑ^+/+^ cells (Figure 2H). Finally, mature CD45.1^+^ peritoneal macrophages did not express RELMɑ after transfer into CD45.2^+^ naive recipients (Figure 2H). Thus, our experimental data support the view that monocytes differentiating into macrophages early after entry into a tissue begin to express RELMɑ independently of IL-4 signalling. These findings add weight to our predicted model of macrophage activation.

### Macrophage activation stages are conserved across tissues and inflammatory conditions

Our findings in the adipose tissue datasets indicate that observed heterogeneity in macrophage activation occurs as these cells enter a tissue and begin transiting through defined activation paths. Our results further indicate that an early stage of activation is characterized by a transient wave of RELMɑ expression, which we confirmed at a distinct site, the peritoneal cavity. Based on this observation, there should be evidence of historical *Retnla* expression in tissue resident macrophages. Consistent with this, historical RELMɑ expression has been reported before in many resident macrophage populations (*35*) and some studies have used RELMɑ expression to identify cells of a distinct tissue-restricted phenotype (*22*) or in an immature state of differentiation (*8*). Our data, which is broadly in agreement with these past reports, indicates that historical RELMɑ expression should be evident in tissues where circulating monocytes replace tissue resident macrophages. Moreover, our data suggest that transit via a RELMɑ^+^ stage is a common feature of all macrophages and not restricted to a single tissue or macrophage subset. To evaluate these ideas further and test to what extent our defined activation paths were conserved across inflammatory conditions and tissues, we set out to use our adipose tissue dataset as a reference to interrogate multiple other situations of tissue inflammation (Supplemental Table 3).

First, we took advantage of a recent data transfer implementation (*36*). This approach identifies pairs of cells across datasets with similar transcriptional profiles, and then uses these “anchors” to transfer data from a reference to a query dataset, assigning a probability of accuracy to the assigned labels. In addition to labels, this approach allows for the imputation of expression data, that is inferring and assigning missing gene expression values, thus enabling the construction of cell atlases despite large technical variations between component data (*36*). Moreover, this strategy was found to be the most accurate tool available (*37*). As anchor selection is critical in this process, we performed extensive benchmarking of the parameters used to find, filter and then apply these transformations, selecting values that would retain only high quality anchors. Moreover, we tested this approach on datasets which contained a mixture of macrophages and other CD45^+^ cells, using the adipose tissue as a reference, reasoning that only macrophages should have high probability scores as a consequence of the activation stage label transfer process (Figure S3A-H). In the first control dataset, where the relative abundance of macrophages to other immune cells was balanced, we observed a bimodal label probability distribution (Figure S3A). Upon closer inspection, we determined that the population of cells with a high label probability could be identified as macrophages, either based on a macrophage gene set score (Figure S3B-C & Supplemental Table 2) or by examining individual macrophage genes (Figure S3D). Using a label probability threshold of 80% (or 0.8) almost exclusively macrophages were assigned a label (Figure S3B-C, colored cells), while most other immune cells were not (Figure S3B-C, gray cells). In a second control dataset, where macrophages made up only a small portion of all CD45^+^ cells (Figure S3E-H), we observed that the label probability distribution was skewed towards 0% (Figure S3E). Nevertheless, applying a similar threshold as before we found that almost exclusively cells with a high macrophage score (Figure S3F-G) and expressing macrophage specific genes (Figure S3H) were labeled. Thus, we felt confident that using the benchmarked parameters in this data transfer approach, as well as the defined threshold, we would be able to interrogate multiple tissues and inflammatory conditions using the adipose tissue dataset as a reference.

We retrieved several publicly available scRNA-seq datasets containing macrophages, representing 9 different tissues and 13 inflammatory conditions with their respective healthy controls, including infections, injuries, cancer and dietary interventions (Figure 3 & Supplemental Table 3). In all cases, we extracted macrophage transcriptomes by calculating a macrophage score as described before, harmonized the data within each tissue to remove batch effects and finally applied the transfer process, assigning cells to the distinct activation stages defined in Figure 1, with a label probability (Figure 3) and imputed gene expression data (Figure S3I-Q). Imputed data was used to cluster and to calculate a UMAP for each dataset. As expected, imputation altered the gene expression values in the original data, however the overall expression patterns were maintained (Figure S3I-Q). We also examined the label probability distribution across identified clusters within each dataset (Figure S4). For any given tissue, we could see that identified clusters would frequently be dominated by a single activation stage label (e.g. Figure S4B, clusters 3, 4, 6 & 7; Figure S4D, clusters 0, 1, 3, 4, 6, 7, 10, 11, 12 & 13), even if not all the cells in that cluster passed the 80% probability threshold previously established. We decided that as all the cells in any given cluster are transcriptionally similar, it was reasonable to assign a dominant label to these cells, even if label probability levels for some of them were below the established threshold (Figure S4). We did this exclusively where a single label was dominant and a sizable portion of the labelled cells passed the confidence threshold. Macrophages that did not meet this criteria were marked as “not classified”, that is cells in clusters where no dominant label was observed or where the label probability was low. Strikingly, our analysis revealed that in all interrogated datasets, we could identify most of the activation stages defined in our reference (Figure 3 - colored cells) with a reasonable proportion of cells with a high label transfer probability (Figure 3).

**Fig. 3.**
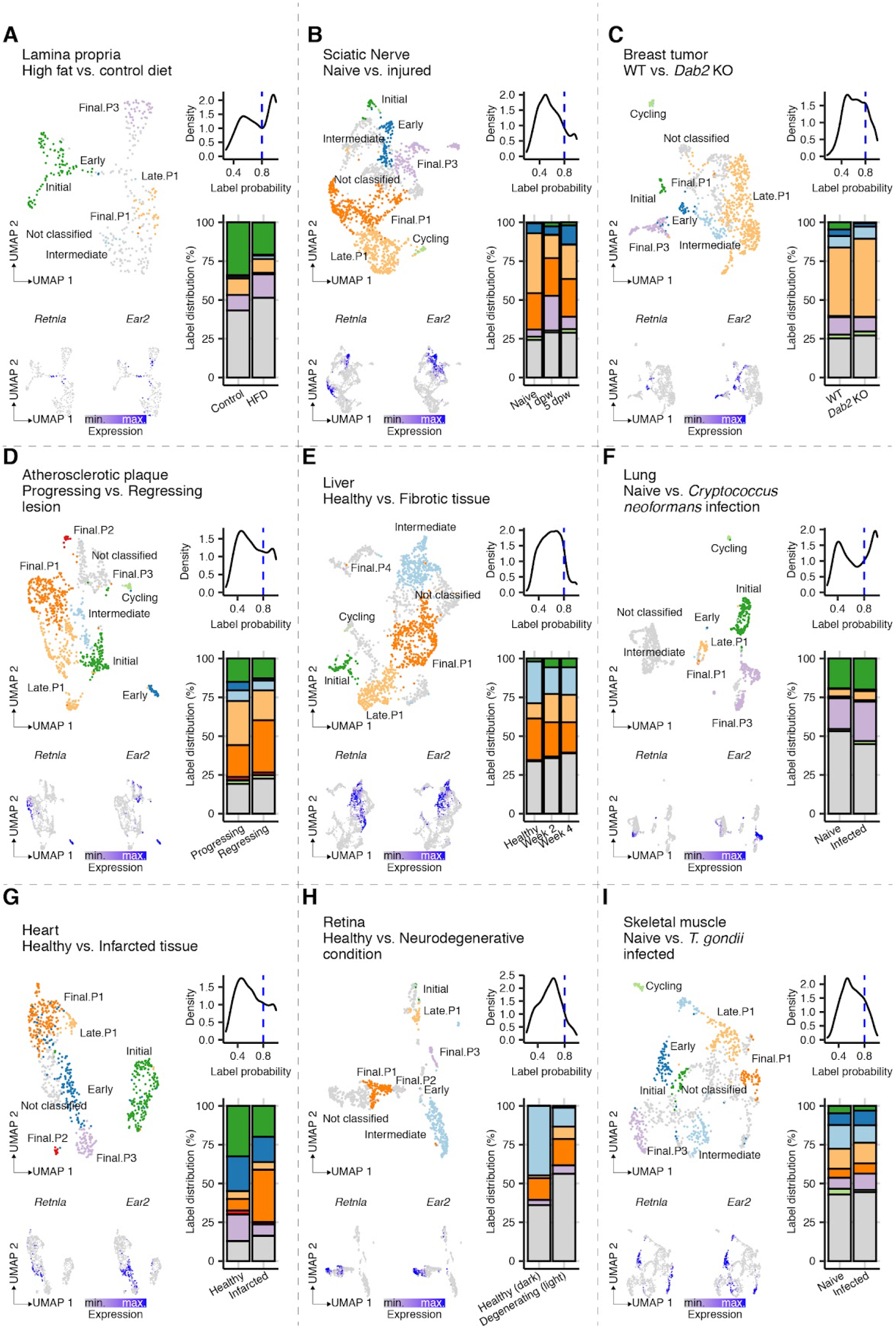
Macrophage activation stages are conserved across tissues and inflammatory conditions. (A-I) Top left - UMAP of macrophages from indicated tissue and condition labelled according to predicted activation stage, including “Not classified” cells (gray). Top right - Label probability distribution from indicated tissue and condition, showing confidence threshold (dashed blue line) for label assignment. Bottom right - Stage distribution shown as a percentage of total cells per biological condition, colored to match predicted labels. Bottom left - UMAP of relative expression (low - gray; high - blue) of *Retnla* and *Ear2*. Cell numbers: Lamina propria - 332; Sciatic nerve - 1500; Breast tumor - 1000; Atherosclerotic plaque - 1000; Liver - 1800; Lung - 1000; Heart - 773; Retina - 897; Skeletal muscle - 1000.

The distribution of activation stage labels was different in each studied tissue and then modified by the corresponding inflammatory conditions (Figure 3). This could reflect the influence of the tissue micro-environment in shaping the immune response, as well as the way in which the immune response is tailored to a specific insult. For example, in the large intestine lamina propria (Figure 3A), infiltrating monocytes in the “Initial” activation stage were abundant in steady state (Figure 3A, bottom right) in line with the reported turn-over of macrophages in this tissue (*9*). However, after 12 weeks of high fat diet (HFD) the proportion of “Initial” stage macrophages diminished, being replaced by cells in the “Final” stage of the “inflammatory” path (Figure 3A, Final.P3 - light purple cells), in accordance with increased inflammation as a result of this intervention (*21*). This correspondence between our labelling strategy and established biology could be seen in all datasets. For instance, in sciatic nerve injury (*22*) a wave of inflammatory cells (Final.P3) could be seen 1 day post wounding (dpw) which receded by day 5, when cells in the “Final” stage of the “phagocytic” path took over (Figure 3B, Final.P1 - dark orange cells). In breast tumors (*23*), the macrophage landscape appeared dominated by cells in the “phagocytic” path (Figure 3C, Late.P1 & Final.P1 - orange cells), which as we mentioned above displayed markers of alternatively activated cells. The same appeared true in regressing atherosclerotic plaque lesions (*24*) (Figure 3D) and in liver fibrosis (*25*) (Figure 3E), while fungal infection in the lung (*26*) resulted in an increase in cells in the “inflammatory” path (Figure 3F, Final.P3 - light purple cells). In infarcted heart (*27*) and retinal damage (*28*), an expansion of cells in the “phagocytic” path was also evident (Figure 3G-H, Late.P1 & Final.P1 - orange cells), although the diversity of activation stages in each tissue was strikingly different, with few identified stages in the Retina both in steady state and after light induced neurodegeneration (Figure 3H). In contrast, skeletal muscle macrophages (*19*) displayed diverse activation stages, with chronic parasitic infection having a modest effect on the stage distribution in this tissue (Figure 3I), although increased “inflammatory” path macrophages were apparent (Final.P3 - light purple cells). Finally, we observed “Early” stage cells, co-expressing *Retnla* and *Ear2*, in nearly all analyzed datasets (Figure 3A-D, G & I), underscoring how this activation step is common to macrophages in most tissues.

Despite demonstrable utility of our labelling approach, in terms of the immediate parallels that could be drawn between the data and published observations, there remained a number of cells with no label assignment (i.e. “not classified”). At least 2 explanations for the abundance of these cells in the studied datasets come to mind. First, our approach hinges on stringently identifying anchor pairs between the data, obtaining a high score and/or having dominant labels in the clusters. Consequently, we are less efficient at identifying transcriptional profiles of cells in between defined activation stages. For this reason, many of the “not classified” cells in our analysis could be seen in between labelled clusters in the UMAP, and would often share low probability labels for flanking clusters with a more defined signature (e.g. Figure S4D, cluster 2 flanked by 1, 4, 7 & 9). Likely for this reason, “Intermediate” stage cells were relatively rare in our analysis of the query datasets, as these were the least defined transitional state that we identified in our reference data. Second, embryonically seeded tissue resident macrophages display a transcriptional profile that is unique (*4, 5*), and thus might not easily relate to activated macrophages originating from circulating monocytes in inflammatory settings. Indeed, we observed the most unclassified cells in tissues where monocyte infiltration is rare (Retina - Figure 3H) or where specialized macrophage subsets are common (alveolar macrophages in the Lung - Figure 3F). In fact, the distinct cluster of “not classified” cells present on the left of the Lung UMAP (Figure 3F) was enriched for alveolar macrophage markers, thus explaining the striking bimodal label distribution in this dataset (Figure 3F). By contrast, the label assignment in the atherosclerotic plaque dataset was nearly global (Figure 3D) likely as only circulating monocytes-derived macrophages were studied (*24*).

We explored the issue of macrophage embryonic origin and tissue immune privilege in more detail by studying a dataset where microglia were recovered and sequenced at different developmental stages from naive mice (Figure 4A and Supplemental Table 3) (*29*). In line with our expectations, we observed very poor label probability distributions for all investigated ages (Figure 4A). Interestingly, the probability threshold was never surpassed and indeed these distributions were skewed progressively towards 0 as the age of the investigated animals increased.

**Fig. 4.**
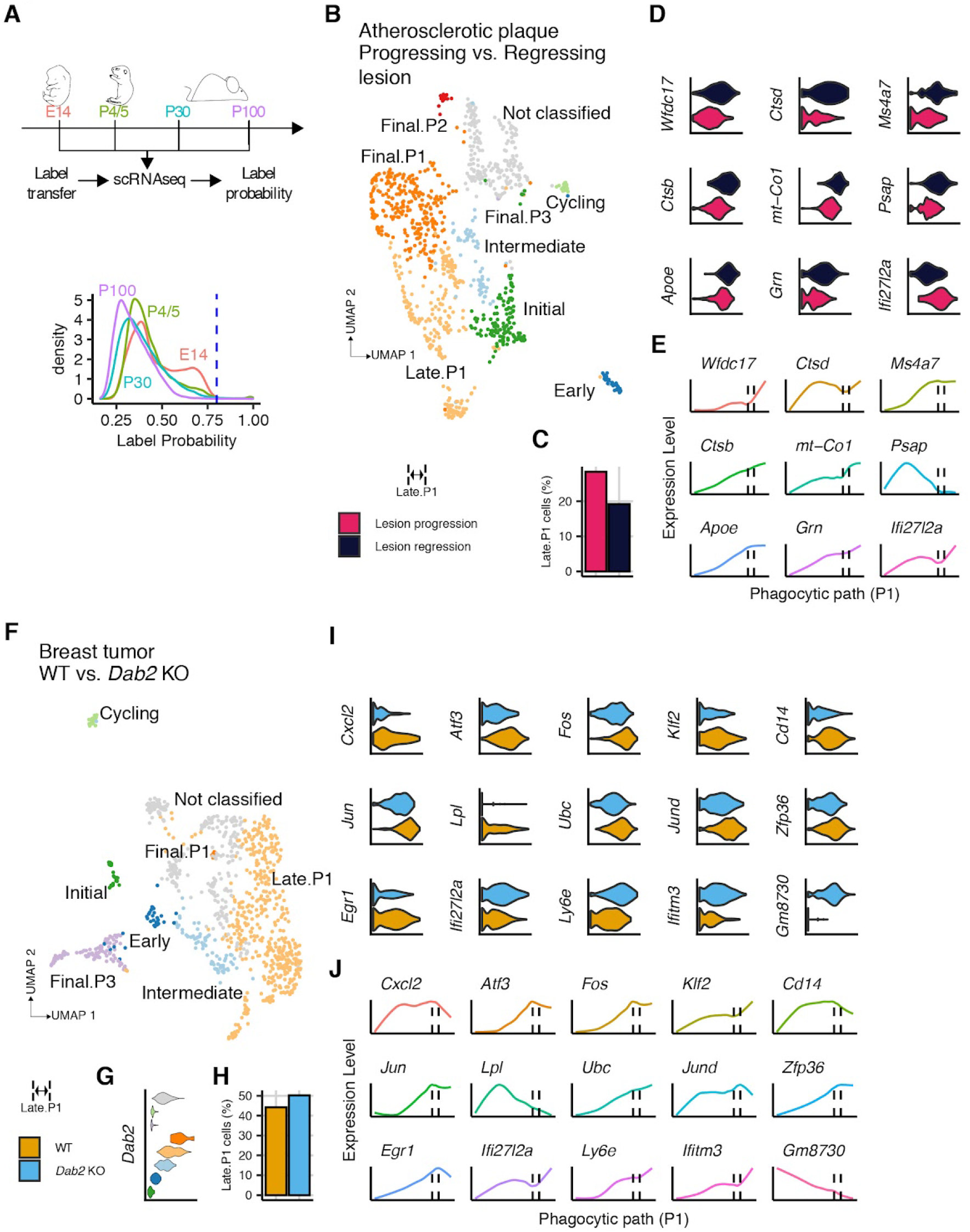
Dysregulation of path-associated gene expression results in pathological macrophage activation stalling. (A) Label probability distribution of microglial datasets obtained from mice at indicated developmental stages. (B) UMAP with stage labels from macrophages obtained from Atherosclerotic plaque regressing (dark blue) and progressing (pink) lesions. (C, H) Percentage of Late.P1 cells per biological condition. (D) Significantly (adjusted p value < 0.01) regulated genes (log fold change > 0.25) in Progressing compared to Regressing lesion macrophages. (E, J) Fitted GAM models for expression of differentially regulated genes (y axis) as a function of Phagocytic path pseudotime (x axis) indicating Late.P1 stage (dashed vertical lines). (F) UMAP with stage labels from macrophages obtained from spontaneous breast cancer tumors in animals with a macrophage specific *Dab2* deletion (*Dab2* KO - light blue) and wild type littermates (WT - ochre). (G) Violin plot of relative *Dab2* expression across activation stages. (I) Significantly (adjusted p value < 0.01) regulated genes (log fold change > 0.25) in *Dab2* KO compared to WT macrophages.

Taken together, our label transfer analysis shows that macrophages across tissues and inflammatory conditions share common transcriptional profiles that correspond to definable activation paths. Our analysis also suggests that embryonically seeded and highly specialized tissue resident macrophages do not respond to inflammatory conditions in a way analogous to that of infiltrating monocytes, with the latter encapsulating most of the functional diversity found in all the studied datasets. Finally, well-established paradigms of macrophage biology are reinforced by the functional stages we defined, making these labels a potential tool to probe deeper into the functional specialization of macrophages during inflammation.

### Exploiting the predictive nature of the proposed macrophage activation model

In light of the predictive nature of our activation model, and its ability to assign activation stage labels to macrophages engaged in inflammatory conditions, we decided to explore in greater detail potential biological insights that might be gleaned from the analysis. For this purpose we turned to the atherosclerotic plaque and breast tumor datasets (Figure 4). In both cases, investigators introduced interventions that ameliorated disease progression (Supplemental Table 3) (*23, 24*). Additionally, we observed in both datasets an alteration in the proportion of macrophages in the “Late” stage of the “phagocytic” path (Figure 3C-D, Late.P1 - light orange). Thus, we reasoned that these data provided an attractive opportunity to explore the activation model more closely.

Dietary and pharmacological intervention (Supplemental Table 3) were reported to induce regression of atherosclerotic plaque lesions (*24*), which are dominated by macrophages in the “phagocytic” path (Figure 4B, dark and light orange cells). Strikingly, there was a shift between “Late” and “Final” activation stages in regressing lesions, with a sizable decrease in the proportion of “Late.P1” cells (Figure 4C). We investigated which genes were altered in expression within cells in this activation stage between progressing and regressing lesions. Critically, we performed this analysis on the original, not on the imputed expression data, guaranteeing that our label predictions served to orient the analysis without affecting the underlying measurements. We then compared these regulated genes with those associated with the “phagocytic” path based on our pseudotime analysis (Figure 2). Interestingly, all but one of the genes in this stage had increased expression in regressing lesions (Figure 4D). Moreover, all but one of these genes tended to be expressed more strongly as cells progressed from “Late.P1” (Figure 4E - dashed lines) to “Final.P1”. Collectively this data suggest that the intervention causing lesions to regress, induced the accelerated transit of macrophages along the “phagocytic” path, as indicated by the increased expression of genes associated with this trajectory in “Late.P1” cells, which concomitantly decrease in abundance.

Next we turned to the breast cancer dataset, where macrophage specific *Dab2* depletion was reported to dampen tumor progression (Figure 4F-J & Supplemental Table 3) (*23*). We observed that *Dab2* was most highly expressed in “Late” and “Final” stage macrophages in the “phagocytic” path (Figure 4F-G), with the former being more abundant. In fact, we observed an increase in “Late” stage *Dab2* deficient macrophages (Figure 4H), leading us to examine differentially expressed genes between these cells and their WT counterparts. As above, we performed this analysis on the original data, using the label assignment only as guidance. Our results show that from 15 regulated genes also included in the pseudotime analysis, 11 were differentially down-regulated between these two groups (Figure 4I). Of these 15 genes, 13 were highly expressed at this stage of activation in our reference data (Figure 4J - dashed lines). The downregulation of these path-associated genes and the accumulation of “Late.P1” cells could indicate that the absence of *Dab2* stalls the progression of macrophages in the “phagocytic” path.

### Monocytes enter wounds populating the functional diversity predicted by the proposed macrophage activation model

Our data indicate that macrophages can be found in similar activation stages in different tissues and conditions, and that the flux of macrophages through these activation stages might be influenced by the immunological processes occurring therein. Moreover, our model predicts that incoming monocytes are able to assume the phenotype of existing macrophages in the tissue and populate all functional stages described. Indeed, in the atherosclerotic plaque dataset, all sequenced cells were derived from circulating precursors (*24*). In order to formally evaluate this, and to validate our predictions, we performed a fate mapping experiment where we traced the influx of monocytes into wounded skin (Figure 5A-B & S5). Red fluorescent monocytes (tdRFP^+^) were administered i.v. 2 or 12 dpw, and all wound macrophages were harvested 4 and 14 dpw (Figure 5B), using index sorting to retain fluorescence values from barcoded cells for further analysis. We mapped, clustered (Figure S5A) and labelled the sequenced cells (Figure 5A) as described above, applying similar thresholds and evaluating label probability distributions for the entire dataset (Figure 5C) and across clusters (Figure S5A-B). Like in other analyzed tissues, wounded skin also exhibited most previously defined activation stages, again demonstrating the robustness of our activation model.

**Fig. 5.**
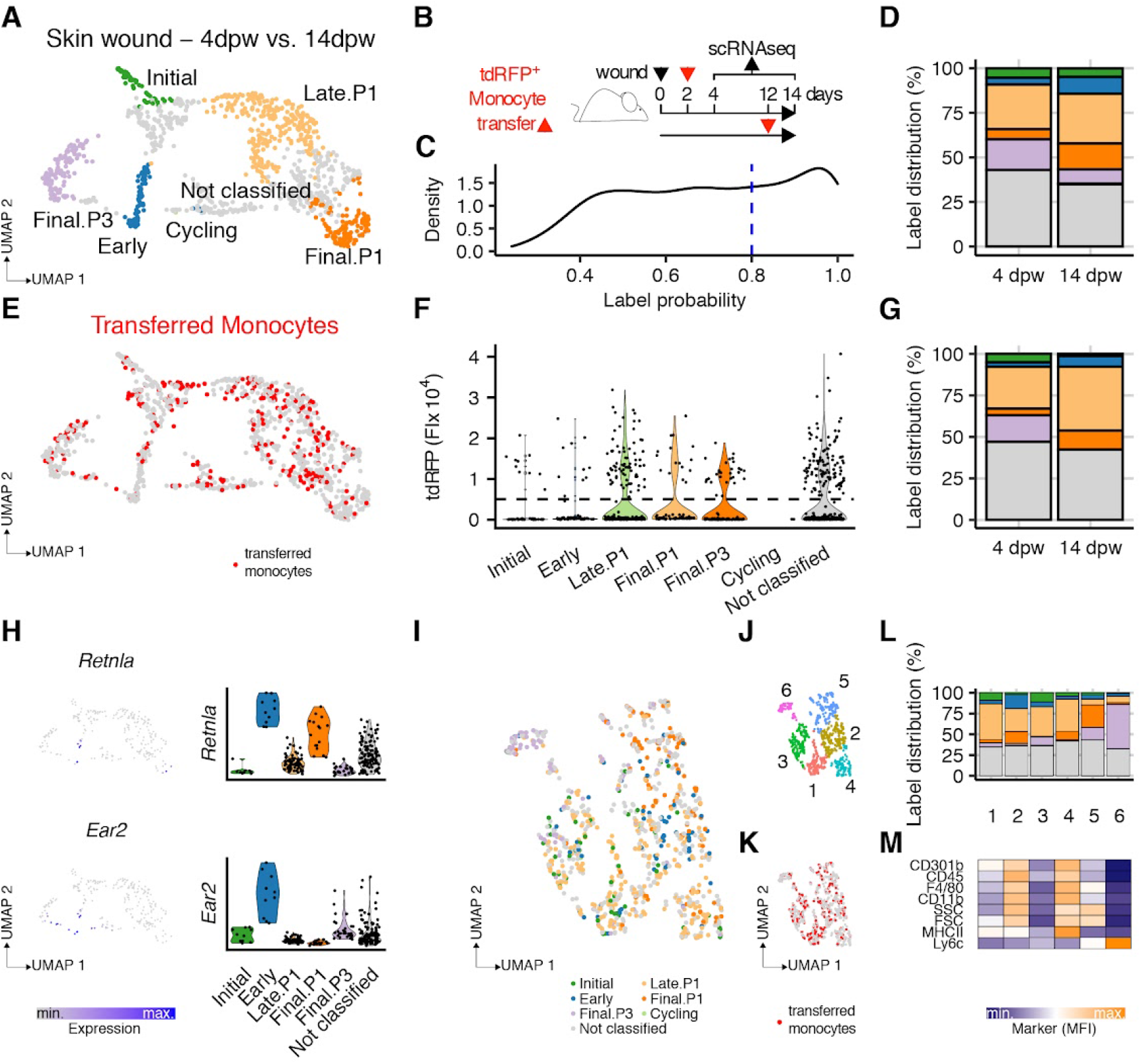
Wound macrophage recruitment confirms activation path model. (A) UMAP with stage labels from macrophages (cells = 1061) obtained from wounded skin biopsies 4 and 14 dpw (n = 5-9). (B) Schematic overview of experimental set up for adoptive tdRFP^+^ monocyte transfer into wounded animals. (C) Label probability distribution showing confidence threshold (dashed blue line) for label assignment. (D) Stage distribution shown as a percentage of total cells per biological condition, colored to match predicted labels. (E) UMAP with tdRFP+ monocytes labelled in red. (F) tdRFP+ fluorescence intensity (FI) in all cells across predicted labels. FI threshold for transferred cell detection is indicated as a dashed line. (G) Stage distribution shown as a percentage of total tdRFP^+^ cells per biological condition, colored to match predicted labels. (H) Relative expression (low - gray; high - blue) in tdRFP^+^ cells of *Retnla* and *Ear2* shown as UMAPs (left) and as violin plots for each activation stage (right). (I-K) UMAP calculated based on indexed flow cytometry data, labelled with predicted activation stages (I), k means clustering (J) or tdRFP+ monocytes (K). (L) Stage distribution shown as a percentage of total cells per flow cytometry cluster, colored to match predicted labels. (M) Relative mean fluorescence intensity (MFI; low - blue; high - orange) in flow cytometry clusters.

We observed a global label probability distribution skewed towards 1 (Figure 5C), indicating a good agreement with our reference data. Moreover, label distribution across conditions was consistent with expectations based on established literature (*38*), with an early wave of inflammatory cells at 4 dpw (Figure 5D, light purple) and a later increase in regulatory “phagocytic” path cells at 14 dpw (Figure 5D, orange). Transferred fluorescent cells were detected at both 4 and 14 dpw (Figure S5C), although only when given on day 2 (Figure S5D). Infiltrating monocytes were distributed across all detected clusters and assigned stage labels (Figure 5E-G), with the distribution in particular mirroring closely the distribution for all macrophages sequenced (Figure 5D & G). As predicted by our model, fluorescent cells were only assigned to the “Initial” stage at 4 dpw (Figure 5G), which is to say 2 days post adoptive transfer. Similarly, only at 4 dpw were fluorescent cells in the “Final” stage of the “inflammatory” path (Figure 5G – light purple) further emphasizing the transitory nature of the early inflammatory wave which occurs during tissue repair. By contrast, transferred monocytes mapped preferentially to the “phagocytic” path at 14 dpw (Figure 5G - orange) and to a lesser extent to the “Early” activation stage (Figure 5G - blue). Collectively, these data support our model, showing that monocytes enter a tissue and flux through distinct activation stages in a dynamic manner, assuming all defined functional stages. The fact that transferred cells only mapped to the “Initial” stage on day 4, and that by day 14 they had assumed other identities, reflects the number of days since these cells accessed the wound.

Our data showed an increase in “Early” stage macrophages at 14 dpw (Figure 5G – blue) compared to day 4, which co-expressed *Retnla* and *Ear2* (Figure 5H - showing transferred cells only). Although we expected to find cells labelled in this manner, as in several other studied tissues, the timing of their appearance is somewhat difficult to explain. Our model would predict that cells in the “Late” and “Final” stages of the “phagocytic” and “inflammatory” paths should have gone through this “Early” phase, yet few cells were labelled as such at 4 dpw (Figure 5G). It is possible that in the 2 days since the adoptive transfer, fluorescent monocytes have indeed gone through this “Early” stage and progressed further in their activation. It is also possible that cells bypass this stage to adapt rapidly during inflammation. Our adoptive transfer data into the peritoneum demonstrated that under steady state conditions 2 days were insufficient to observe *Retnla* expression in infiltrating cells (Figure 2G). In fact, the increased proportion of “Early” stage cells at 14 dpw (Figure 5G), is in line with our earlier findings that after 8 days a higher percentage of infiltrating cells would fall in this stage. Thus, we propose that the relative speed at which macrophages traverse defined activation paths is influenced by the inflammatory conditions at the site of immunological insult, so that cells may accelerate their passage through identified stages to better adapt to the required immune response.

Next, we took advantage of the indexed nature of the skin wound dataset to validate our labelling strategy with common macrophage phenotypic markers. For this purpose, we scaled the fluorescent signal of CD301b, CD45, F4/80, CD11b, MHCII and Ly6C, as well as the side and forward scatter parameters, across all cells and calculated a UMAP for this flow cytometry data, retaining the stage labelling based on the transcriptional profile of the cells (Figure 5I). We then identified clusters using k-means (Figure 5J & S5E). Strikingly, we observed that cells in the “Late.P1”, “Final.P1 and “Final.P3” stages each dominated a cluster (Figure 5L - clusters 4, 5 and 6 respectively), while “Early” stage cells localized predominantly to cluster 2 (Figure 5J&L). As expected, transferred monocytes were evenly distributed across these clusters (Figure 5K & S5F), further emphasizing the ability of these cells to differentiate into all identified functional stages. Finally, we show that these flow cytometry based clusters are associated with significantly different protein expression levels (Figure 5M - p value < 0.001).

Collectively, these data show that monocytes entering a wound respond to this environment by becoming activated in accordance to the model proposed in this study. These cells gained access to the inflamed tissue and followed distinct activation paths towards different functional outcomes. The relative proportion of stage labels our model assigns to these cells, mirrors the expectations of established wound repair paradigms, while also validating the observations we have made in other studied tissues. Overall, our results show that proposed activation stages are not only distinct in their transcriptional profile, but that they can be observed based on protein expression.

### Macrophage activation stages in tissues have distinct transcriptional markers

Our approach to define activation stages and paths has relied so far on the use of a reference to interrogate each queried dataset. This approach revealed a striking conservation of macrophage activation dynamics, despite differences in tissue of origin and inflammatory condition. We detected all reference activation stages in one or more tissues (Figure 6A), and we discussed how the relative abundance of each activation stage and the presence of “not classified” cells may be explained. However, our approach relied on the quality of anchor pairs identified across datasets to transfer labels and impute gene expression data. As these processes by necessity transform the original expression data, we sought to determine the robustness of our approach by interrogating labelled cells directly (Figure 6B). For this purpose we took all high label probability cells (>80%) from 10 query datasets, and combined these with a randomly sampled portion (n= 500) of macrophages from our reference, retaining only the label assignment and original uncorrected gene expression data (Figure 6B). Once extracted, these macrophages (n = 2843) were integrated across tissues, without giving priority to any dataset, then clustered and visualized as a UMAP (Figure 6C).

**Fig. 6.**
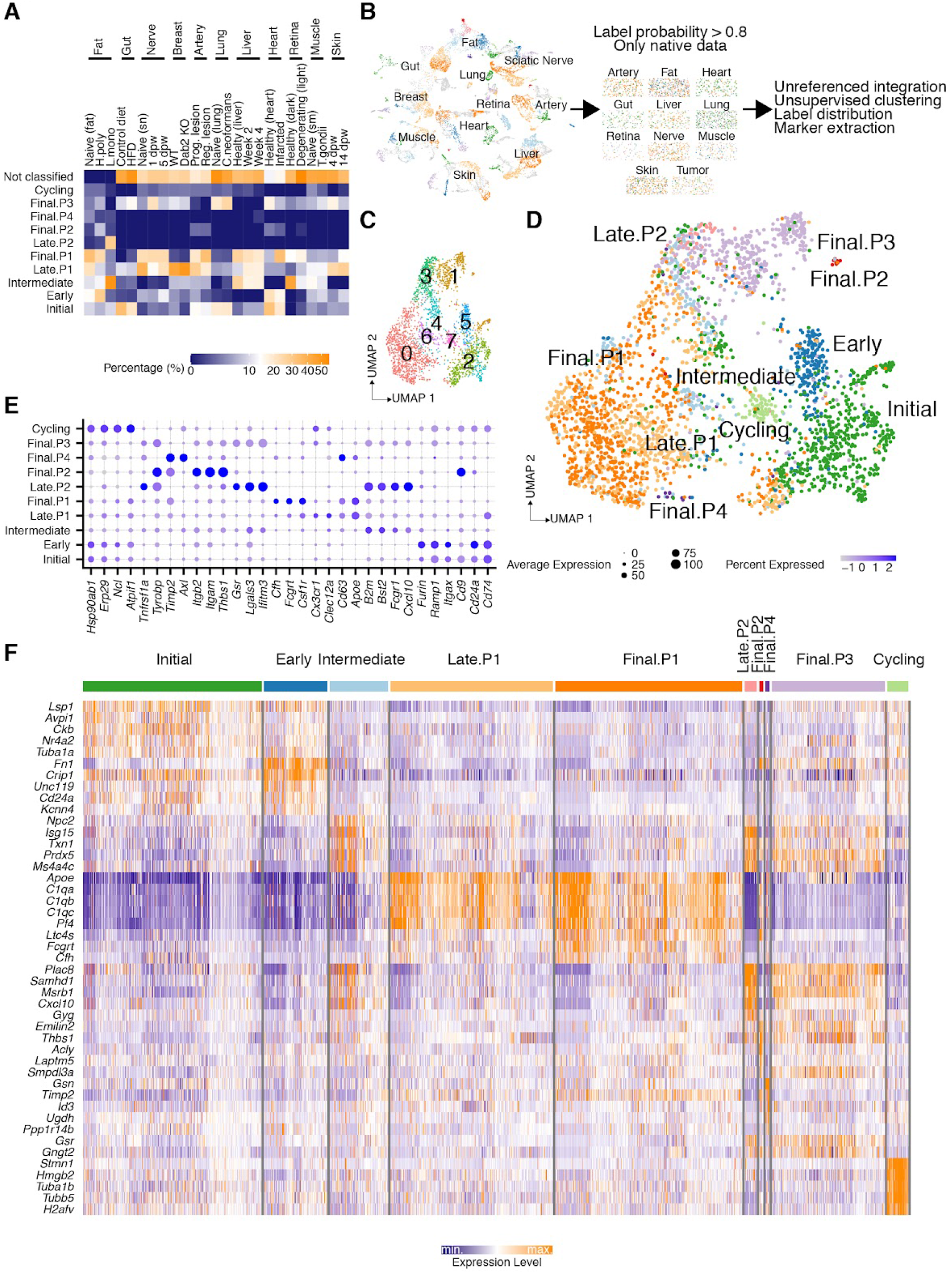
Cross-condition data integration reveals stage-specific marker genes. (A) Activation stage distribution shown as percentages of total cells per biological condition and tissue. Scale was modified with a square-root transformation for ease of visualization. (B) Schematic overview of data reprocessing strategy. (C-D) Integrated UMAP of macrophages (cells = 2843) across conditions and tissues labelled with identified clusters (C) or previously assigned activation stages (D). (E) Dot plot of top significantly regulated genes associated with GO term “Cell surface” (GO:0009986) across activation stages. (F) Heatmap showing relative expression (low - blue; high - orange) of top significantly (adjusted p value < 0.01) regulated genes (log fold change > 0.25) across activation stages.

Our expectation for this analysis was that the transcriptional tissue signature would not obscure the activation stage label. That is, our model would predict that tissues would not define the resulting cell clustering, but rather that the activation stage of these cells would be sufficient to group them in this unsupervised analysis. Critically, this was the outcome we observed (Figure 6D). Cells with identical activation labels clustered together, regardless of the tissue of origin or the inflammatory condition. This demonstrated that our initial anchor, label transfer and data imputation approach was valid. Importantly, having all macrophages clustered in this manner, allowed for the extraction of tissue-independent transcriptional markers for all activation stages. Indeed, we found genes associated with cell surface expression (Figure 6E & Supplemental Table 5) and other upregulated genes (Figure 6F & Supplemental Table 4) corresponding to each identified activation stage, which largely aligned with the genes we originally associated with each label (Figure S1C & Supplemental Table 1).

Finally, we observed that integrated macrophages were organized similarly to our reference data. The “Initial”, “Early” and “Cycling” stages clustered near each other, while the “Intermediate” stage separated the “phagocytic” and “remodelling” paths at the bottom from the “oxidative stress” and “inflammatory” paths at the top of the UMAP (Figure 6D). This distribution of clusters adds weight to our proposed activation trajectories and emphasizes the relative relationship between the activation stages defined.

### Transcriptional network analysis reveals macrophage gene expression hubs and upstream regulators

Our analysis has shown that macrophages in inflamed tissues flux through conserved activation paths in order to respond to inflammatory insults. Similarly, we have established that these activation paths and stages are robust and associated with distinct transcriptional profiles that provide functional insights about these cells. Lastly, we hypothesized that changes in the regulation of genes associated with defined activation paths was likely to stall or promote macrophage activation. To explore this final aspect of the data more closely, we returned to the genes that we associated with pseudotime (Figure 2) and built a network based on known protein-protein interactions (Figure S6A), calculating the edge weight as a combination of the confidence of the interaction and the goodness of the model fit across all paths for the pair of connected nodes. We then filtered the network, to retain only high weight edges (Figure S6B, edge weight threshold blue dashed line) and removed disconnected nodes. Finally, we clustered the resulting network, performed GO term enrichment analysis in each cluster (Figure S6C) and labelled the network according to the most enriched term in the gene set. The resulting network (Figure 7A, 242 genes) shows 10 clusters associated with “Superoxide metabolic process”, “Leukotriene synthesis”, “Response to external stimulus”, “Lipid synthesis”, “Myeloid differentiation”, “Antigen processing and presentation”, “Protein complex oligomerization”, “Chemotaxis”, “Lipid transport” and “Cytoskeleton organization”. These gene sets are not engaged similarly by all activation paths, as revealed by subsetting the network to include only genes significantly associated with pseudotime in each path (Figure 7B).

**Fig. 7.**
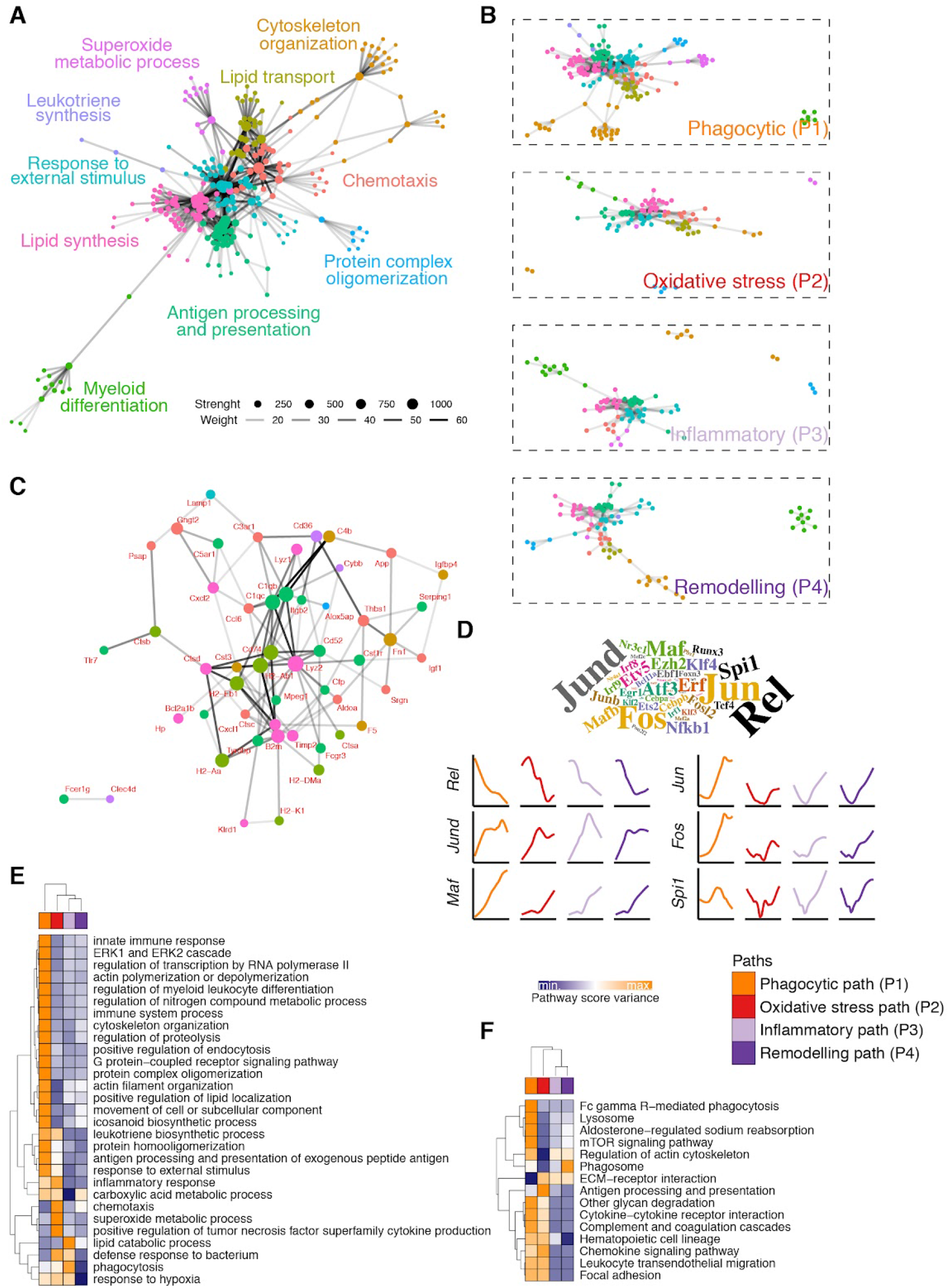
Macrophage transcriptional network across activation paths. (A) Transcriptional network of protein-protein interactions, depicting genes (n = 242) as nodes and interactions as edges (n = 716). Node size corresponds to calculated strength. Node color is associated with the assigned network cluster. Edge opacity corresponds to calculated weight of interaction. The most enriched GO term associated with each cluster is indicated. (B) Transcriptional network as in A, but split along activation paths, with arbitrary node size and edge opacity. (C) Transcriptional network as in A, limited to nodes with high betweenness (75% quantile) that connect 2 or more clusters. Gene names indicated in red. (D) Top - Transcription factor enrichment analysis shown as a word cloud where size of name is proportional to the number of gene sets in transcriptional network clusters associated with the specific transcription factor. Bottom - Fitted GAM models (colored lines matching activation paths) of gene expression of enriched transcription factors (y axis - fixed across all paths for each gene) as a function of pseudotime (x axis - specific to each path). (E-F) Heatmap showing relative gene set score variance (low - blue; high - orange) of enriched GO terms (E) or KEGG pathways (F).

We next extracted three types of information from this transcriptional network. First, we examined which genes had the potential to act as central nodes of information transfer, as these could be targets for therapeutic intervention. We reasoned that these information hubs could be represented by highly connected nodes, which articulated the network by connecting 2 or more clusters, and that were overrepresented in the paths connecting pairs of nodes in the network (i.e. high betweenness). Examining genes meeting these criteria (Figure 6C) highlighted both some well-known macrophage regulators (e.g. *Lyz2*, *Csf1r*), but also genes whose function has not been widely studied in the context of macrophage activation (e.g. *Gngt2*, *Srgn*), thus warranting further exploration. Second, we identified transcription factors (TF) upstream of the transcriptional network clusters, which were themselves regulated dynamically along the activation paths. We ranked these based on the number of times they were associated with a gene set in our clusters (Figure 7D, word cloud) and show the dynamic regulation of 6 TF in each activation path (Figure 7D, bottom). Interestingly, some TF had opposing behaviours (*Rel* vs. *Maf*), while others behaved similarly in all paths except one, where they suddenly veered in opposing directions (*Spi1* vs. *Fos*/*Jun*). As these sudden direction changes, as well as the relative levels of one TF to another could represent important decision points in activation paths, these TFs too warrant further examination. Finally, we wished to provide a more detailed process enrichment profile for each activation path. Consequently, we took the top 3 GO enriched terms in each cluster (Figure 7E & S6C) and all enriched KEGG pathways detected (Figure 7F) and calculated gene set scores for each of these (Supplemental Table 2). We then estimated the variance of every gene set score within cells of each activation path and represented these data as a heatmap (Figure 7E-F), finding a distinct profile for each path for these functions. In this manner we highlight the relative regulation of several processes of interest in the activation paths we defined, as a guide to researchers wishing to explore these aspects of macrophage function in more detail.

In summary, we employed a predictive model of label transfer to encompass all forms of macrophage activation irrespective of tissue or inflammatory condition. We demonstrate that this model is robust, aligning with well-established paradigms of macrophage function, while providing novel avenues for investigation. We provide surface and global gene expression profiles for these activation stages to aid in their identification in future studies. Lastly, we have prepared an online tool (https://www.macrophage-framework.jhmi.edu) to aid in exploring the data contained in this study. Our results emphasize the conservation and relative homogeneity of macrophage activation across tissues, transcending macrophage tissue residence, while still allowing for activation diversity.

## Discussion

Advances in the understanding of macrophage ontogeny and of differential gene expression signatures linked to macrophage tissue residence has revealed inherent complexity within this cell type. Moreover, the transcriptional profile of macrophages following exposure to a broad range of stimuli for which they are known to express receptors revealed a spectrum of potential activation states not captured by *in vitro* models. In light of these findings, it has become difficult to relate macrophage activation across investigations. Our study offers an alternative view of macrophage activation in tissues during inflammation. By comparing the transcriptional profiles of macrophages recovered from different tissues from mice experiencing distinct diseases/conditions, we identified a limited and consistent number of transcriptional profiles that were unobscured by the tissue or stimulus studied. We modelled these conserved and yet diverse signatures as stages across four activation paths, finding that “phagocytic” and “inflammatory” paths were most common. These paths have features in common with M2 and M1 respectively, encompassing those references while offering a broader and dynamic alternative. Finally, our analysis offers insights into the information hubs, transcription factors and gene expression programs that are responsible for shaping macrophage function.

The macrophage activation model we propose, where cells transit through “initial” and “early” stages of commitment to a particular path, is evident in other independent analyses. For instance, in a murine model of non-alcoholic steatohepatitis (NASH) a monocyte derived population of Ly6C^lo^ macrophages expressed high levels of *Ccr2*, *Klrd1* and MHC-II (*39*), comparable to genes expressed in “initial” stage cells in our analysis (Supplemental Table 1 and https://www.macrophage-framework.jhmi.edu). *Ccr2* expression in particular gives credence to our choice of starting point for the model as this encodes a critical tissue-homing receptor in circulating monocytes (*40*). Notably, a closely associated cell population in the NASH dataset expressed both *Ear2* and *Fn1*, mirroring “early” stage macrophages (Supplemental Table 1).

Moreover, in the context of this disease, this population gave rise to Kupffer cells (*39*), which expressed high levels of *Mrc1*, *Apoe* and complement associated genes, similarly to “phagocytic” path macrophages. Another instance where this progression is evident is in joint synovial macrophages (*41*). In this setting, two populations of interstitial macrophages, one MHC-II^high^ and one RELMɑ^+^ reminiscent of the “initial” and “early” stages described herein, respectively, replenished long-lived synovial tissue-resident cells (*41*). The possibility to reconstruct our model in these independent analyses demonstrates the robustness and universality of our findings.

Our model highlights the role of incoming monocytes into tissues, both under homeostasis and inflammatory conditions. The input of monocyte-derived macrophages to the overall tissue macrophage pool during homeostasis varies from organ to organ (*4*), and under these conditions we found that the contribution of the four identified activation paths was intriguingly diverse between tissues, likely as a result of microenvironmental signals that are themselves heterogenous. Thus, our data indicate that the commitment of monocytes to these activation paths is regulated not just by the inflammatory settings, which invariably altered the proportion of cells in each stage, but also the specific nature of the tissue. We further demonstrated that the emphasis of our model on monocyte-derived macrophages was likely due to the unique transcriptional profile of embryonically derived macrophages (*2*), which were labeled as “not classified” in our analysis. One implication of this finding is that incoming monocytes give rise to most of the functional diversity in any given tissue, while the resident cells remain more transcriptionally stable regardless of the insult. Similar conclusions were drawn recently (*10*) from observations on alveolar macrophages, which have been shown to be less plastic, less phagocytic, more permissive to infection, less responsive to IL-4 stimulation, and generally less engaged in ongoing local immune responses than are monocyte-derived cells (*42–44*). Tissue resident macrophages in other tissues, specifically the peritoneal cavity, have also been shown to be less immunologically active (*45*), even if they are highly proliferative (*46*). Likewise, monocyte-derived macrophages have been shown to play a dominant role in tumors (*47*). The mechanisms restricting tissue-resident macrophage activation have not been elucidated, although epigenetic imprinting (*1, 10*) and autophagy-enforced quiescence (*48*) are likely candidates. Overall, the emerging picture is one where macrophage functional plasticity in response to a loss of homeostasis within tissues, is a feature of cells derived from recruited monocytes, which participate in the induction and resolution of inflammation by moving along defined activation paths. This view does not exclude the possibility that tissue resident macrophages are contributing to the response to tissue damage, but it does predict that monocyte-derived cells are the major contributors in this regard.

In our model, we postulate that macrophages become activated through 4 possible paths and that these paths are unidirectional such that cells become committed exclusively to one route. We infer that only cells in the “phagocytic” path go on to replace tissue-resident macrophages. Several independent lines of investigation support this hypothesis: Increased expression of complement genes has been reported in Kupffer cells (*39*) and alveolar macrophages (*49*) derived from monocytes; Phagocytosis appears to be a key feature of tissue resident macrophages (*50*); Phagocytic receptors like *Mrc1*, *Cd163*, *Timd4* and *Mertk*, all highly expressed in “phagocytic” path cells, are associated with tissue resident macrophages (*50*). It is possible that through phagocytosis, macrophages become tissue imprinted. Thus, by engulfing apoptotic cells, macrophages might indirectly absorb factors that convey tissue identity.

By far the most abundant gene signature we observed in our analysis was that of the later stages of the “phagocytic” path. As mentioned above, this transcriptional profile is evident elsewhere (*39, 49*). Indeed, during lung fibrosis *Apoe* and complement gene expression became dominant features of disease progression (*20*). Interestingly, this gene signature can be extended to human macrophages involved in injury resolution (*51*). Our proposal that these cells give rise to tissue-resident macrophages explains in part this relative abundance. However, their transcriptional profile also overlaps with genes associated with alternative activation (*17*).

Moreover, the complement product *C1q* has been linked to macrophage proliferation (*52*), a characteristic of alternatively activated macrophages. It is intriguing that these macrophages are critical to restoring homeostasis by removing dead cells, which boosts their IL-4 driven phenotype (*53*), yet they also have a clear role in the pathology of several of the conditions explored herein. Indeed, our findings suggest that stalling macrophages along the “phagocytic” path can be both beneficial, as seen in the case of breast tumors, or detrimental, as observed in atherosclerotic plaques (Figure 4). In fact, even tissue-imprinted pathogenic microglia associated with Alzheimer’s disease converged into an *Apoe*-expressing phenotype (*54*). Based on this, we postulate that determining how to manipulate progression along the “phagocytic” path may offer therapeutic opportunities.

While in depth identification of the drivers of macrophage transit through the proposed activation paths is beyond the scope of this study, our initial exploration of the data revealed 242 highly interconnected genes with some common upstream regulators and mapping to diverse functions. How this transcriptional network is shaped within each tissue will be of great interest moving forward. However, the fact that the number of macrophage activation stages we defined was conserved and limited, despite the diversity of insults and tissues studied, suggests that common undercurrents guiding macrophage activation might be built into tissues. One potential set of candidates for orchestrating these processes would be signals associated with tissue damage, which is ubiquitous during inflammation. Indeed, the production of alarmins by stromal cells leading to the activation of resident innate immune cells (e.g. innate lymphoid cells, mast cells) might be an important driver of macrophage tissue engraftment, particularly as the signals they produce are capable of guiding both pro- and anti-inflammatory phenotypes. Thus, seemingly opposing signals (i.e. IL-4, TNF, IL-1β, IL-10, IL-13, PGE2) produced concomitantly by tissue-embedded mast cells (*55, 56*) and ILC2s (*57*) might be partly responsible for the diversity of observed macrophage activation paths in all conditions. It is feasible that additional signals provided by metabolites (*58*) might contribute to these outcomes.

Especially as we move into the era of single cell genomics, establishing a lingua franca that allows us to describe macrophage biology in humans and other animals, and across tissues and diseases, is critical. Not only is this a matter of transferring insights from one study to another, but also in shaping our understanding of the function of macrophages *in vivo*, especially in inflammatory diseases. Moreover, moving the focus away from individual genes and towards gene signatures, might allow for better transferability of findings between mouse and human models. Indeed, our data readily finds parallels in human conditions (*51*). Finally, understanding how the local microenvironment shapes the immune response is possible only if we are able to define the common threads of that response in the first place. In this context, our approach highlights the overarching similarity that can be found in the way in which macrophages diversify their function, without dismissing the influence that inflammatory conditions and tissue niches impose on that functionality. We consider our approach is a step towards building a common framework to describe macrophage activation that can be applied broadly to explore the biology of these important cells.

## Materials and Methods

### Data and Code Availability

Publicly available R packages were used to analyse all data contained within this manuscript. Relevant packages are referenced through-out the methods section. Annotated code to reproduce key analysis modules and recreate figures was deposited on github and may be accessed at https://github.com/davidsanin/Macrophage_framework.

An interactive shiny application to visualize critical aspects of the data in this publication can be accessed at https://www.macrophage-framework.jhmi.edu. scRNAseq datasets generated specifically for this publication may be retrieved from publicly available repositories with no restrictions on their use, under the following accession numbers: helminth infection of adipose tissue (GSE157313), bacterial infection of adipose tissue (GSE171328), High fat diet lamina propria (GSE171330) and skin wound (GSE). Accession numbers, associated publications and experimental details of these and all other datasets included in this study are listed in Supplemental Table 3.

### Mouse Models

C57BL/6J (RRID: IMSR_JAX:000664), B6.129P2-Lyz2^tm1(cre)lfo/J^ (RRID: MGI:5014089) and CD45.1 congenic (RRID: IMSR_JAX:002014) mouse strains were purchased from The Jackson Laboratory. Mice were macrophages were deficient for the expression of IL-4Rɑ (IL-4Rɑ^-/-^) were generated by crossing B6.129P2-Lyz2^tm1(cre)lfo/J^ with B6-Il4ra^tm(loxp)^ (*59*). These strains were maintained at the Max Planck Institute for Immunobiology and Epigenetics. Experimental procedures including helminth or bacterial infection, adoptive cell transfer into the peritoneum and experimental diets were performed at the Max Planck Institute for Immunobiology and Epigenetics. Animal care was undertaken in accordance with Institutional Animal Use and Care Guidelines with approval by the animal care committee of the Regierungspraesidium Freiburg, Germany. All animals used for tissue harvest or experimental procedures were female and aged between 6-8 weeks at the start of the experiment. Animals were humanely sacrificed by carbon dioxide asphyxiation followed by cervical dislocation and tissue dissection. Mice were bred under specific pathogen free standards.

For skin wound model, B6.RFP mice with ubiquitous tdRFP expression were generated by germline excision of the loxP flanked STOP cassette (LSL) in R26^LSL-tdRFP^ animals (*60*) employing the pgk-Cre transgene (*61*). This strain, alongside recipient C57BL/6JRj (RRID: MGI:2670020) mice were housed in individually ventilated cages under specific-pathogen free environment at the Experimental Center of the Medical Faculty, TU Dresden. Wound and adoptive cell transfer experiments were conducted according to institutional guidelines and in accordance with the German Law for Protection of Animals approved by Landesdirektion Dresden (TVV 62/2015).

### Adipose tissue infection and cell isolation

#### Experimental infections

L3 infectious stage *Heligmosomoides polygyrus* (*H. poly*) larvae were kindly provided by Dr. Joseph Urban Jr, USDA, ARS, Beltsville Human Nutrition Research Center, Diet Genomics and Immunology Laboratory, Beltsville, USA and maintained at 4°C until required. To induce *H. poly* infection mice were gavaged with 200 L3 infectious stage larvae in PBS. Mice were left for 13 days before being sacrificed.

A wild type strain of *Listeria monocytogenes* (*L. mono*) was used for infections. Mice were infected subcutaneously on the footpad with a sublethal dose of 1 × 10^6^ colony-forming units (CFU). Mice were left for 1 day before being sacrificed.

#### Stromal vascular fraction isolation

For isolation of cells from mesenteric adipose tissue, mice were euthanized and transcardially perfused with ice-cold PBS. Adipose tissue was separated from lymph nodes and surrounding organs (i.e. intestine, omentum), minced and digested in low glucose DMEM (Gibco) containing 25 mM HEPES, 1% low fatty acid bovine serum albumin, 2 mM L-glutamine, 100 U/mL Penicillin/Streptomycin, 0.2 mg/mL Liberase TL (Roche) and 0.25 mg/mL DNase I (Roche) for 30-40 min at 37°C with gentle rotation. After digestion DMEM containing 2 mM EDTA was added and suspension filtered through a 70 µm strainer. Cells in stromal vascular fraction (SVF) were separated from the adipocyte layer by centrifugation.

For isolation of cells from popliteal adipose tissue, mice were euthanized and adipose tissue was separated from lymph nodes and surrounding muscle. Isolated tissue was then minced and digested in DMEM (Gibco) containing 2.5% bovine serum albumin, 2 mM L-glutamine, 100 U/mL Penicillin/Streptomycin, 2 mg/mL Collagenase I (Thermo) and 2 mg/mL Collagenase II (Thermo) for 45 min at 37°C with gentle rotation. During the last 15 minutes of incubation 2 mM EDTA was added to the media. Finally, the cell suspension was filtered through a 100 µm strainer. Cells in stromal vascular fraction (SVF) were separated from the adipocyte layer by centrifugation.

Single live cells were further purified via fluorescent activated cell sorting, excluding dead cells labelled with LIVE/DEAD Fixable Aqua Dead Cell Stain Kit and doublets based on side and forward light scatter.

#### Single cell barcoding and library preparation

scRNA-seq of SVF cells was performed using a 10X Genomics Chromium Controller. Single cells were processed with GemCode Single Cell Platform using GemCode Gel Beads, Chip and Library Kits (v2) following the manufacturer’s protocol loading sufficient cells to obtain 5000 cells per lane following manufacturer’s guidelines. Libraries were sequenced on HiSeq 3000 (Illumina) to achieve 50000 reads per cell.

### Macrophage isolation and adoptive cell transfer

#### Monocyte isolation and transfer

A single cell suspension from bone marrow isolated from freshly sacrificed CD45.1^+^ or IL-4Rɑ^-/-^ mice was prepared by flushing the the tibia and femurs of dissected animals with PBS, then passing resulting suspension through a 70 µm strainer. Red blood cells were removed by brief incubation in ACK Lysing Buffer (ThermoFisher scientific). Cell suspension was incubated in 1% fetal bovine serum in PBS for 30 min on ice with fluorochrome-conjugate monoclonal antibodies plus anti-CD16/CD32 (Biozol) and finally dead cells were labelled with LIVE/DEAD Fixable Aqua Dead Cell Stain Kit (Thermo scientific) following the manufacturer’s instructions. After staining, bone marrow cells were maintained in 1% fetal bovine serum in PBS at 4°C, then target population isolated using a BD FACSaria^TM^ Fusion cell sorter into 50% fetal bovine serum in PBS. Used fluorochrome-conjugate monoclonal antibodies included: CD11b (Biolegend, clone: M1/70), F4/80 (Biozol, clone: BM8), SiglecF (BD Horizon, clone: E50-2440), Ly6G (BioLegend, clone: 1A8), CD11c (BioLegend, clone: N418), MHC-II (BioLegend, clone: M5/114.15.2), Ly6C (BioLegend, clone: HK1.4).

Sorted bone marrow monocytes were stained with Cell Trace Violet (Life Technologies), following the manufacturer’s instructions, counted and then 5x10^5^ cells/mouse were transferred via intraperitoneal injection to littermates randomly assigned to experimental groups.

#### Peritoneal macrophage isolation and transfer

Peritoneal lavage was harvested from CD45.1^+^ by injecting 10 mL of ice-cold 2% fetal bovine serum in PBS into the peritoneal cavity of freshly sacrificed animals, then gently tapping the sides of the mouse to dislodge peritoneal cells, followed by slow retrieval of lavage solution. Cells were recovered via centrifugation and stained as described above using a biotinylated monoclonal antibody against TIM4 (Miltenyi Biotec, clone: REA999), the isolated with anti-biotin Microbeads (Miltenyi Biotec, Cat# 130-090-485) following the manufacturer’s instructions.

Purified TIM4^+^ tissue resident macrophages were then stained with Cell Trace Violet (Life Technologies), following the manufacturer’s instructions, counted and then 5x10^5^ cells/mouse were transferred via intraperitoneal injection to littermates randomly assigned to experimental groups.

### Monocyte recovery and RELMα staining

Peritoneal lavage was harvested from mice 2, 4 and 8 days post adoptive cell transfer, by injecting 10 mL of ice-cold 2% fetal bovine serum in PBS into the peritoneal cavity of freshly sacrificed animals, then gently tapping the sides of the mouse to dislodge peritoneal cells, followed by slow retrieval of lavage solution. Cells were recovered via centrifugation and stained for flow cytometric analysis as described above using the following fluorochrome-conjugate monoclonal antibodies: TIM4 (BioLegend, clone: F31-5G3), CD45.1 (BioLegend, clone: A20) and CD115 (BioLegend, clone: AFS98). Detection of intracellular *Retnla* mRNA and RELMα protein was performed using PrimeFlow RNA Assays (Thermo scientific) following the manufacturer’s instructions. Briefly, surface-stained peritoneal lavage cells were fixed and permeabilized using the kit’s reagents, then incubated with anti-RELMα primary antibody (Peprotech, Cat# 500-P214), and subsequently an anti-rabbit secondary antibody (Life technologies). Probe hybridization, signal amplification and fluorochrome conjugation with PrimeFlow target-specific probes for *Retnla* and *Actb* (as a positive control) were carried out over the course of 2 days following the manufacturer’s guidelines. Data from stained cells were collected using LSR Fortessa flow cytometers (BDBiosciences) with FACSDiva v. 9.0 and data were processed using FlowJo v. 10.6.

### Dietary intervention and large intestine lamina propria cell isolation

#### High fat diet treatment

Obesity was induced by ad libitum feeding of C57/BL6 mice for 12 week with irradiated high fat diet (Rodent Diet 60% kcal from fat, Research Diets, Inc., Cat #D12492). Control diet (chow) containing 24% protein, 47.5% carbohydrate, and 4.9% fat, was given to age and sex matched animals as a control group.

#### Cell isolation

At the end of treatment, animals were sacrificed, and cells from large intestine lamina propria recovered. Briefly, the large intestine was separated at the junction with the cecum, and all remaining connective and fat tissue removed. Isolated intestine was then opened longitudinally, cleaned, cut into 0.4-1 cm pieces and washed with ice-cold 25 mM HEPES in PBS. Tissue fragments were then placed in RPMI (Gibco) medium containing 3% fetal bovine serum, 25 mM HEPES, 5 mM EDTA plus 3.5 mM Dithiothreitol and incubated for 15 min in 37°C with gentle agitation. Tissue fragments were recovered via filtering and vigorously washed thrice with 2 mM EDTA in RPMI, discarding supernatants. Washed fragments were minced then incubated in RPMI supplemented with 0.5% fetal bovine serum, 2 mg/mL Collagenase VIII (Roche) and 0.5 mg/mL DNAse I (Roche) for 30-40 min at 37°C with gentle agitation. Cell suspension and tissue fragments were filtered through a 70 μm strainer, and dissociated with the rubber end of a syringe plunger. Resulting cell suspension was centrifuged and further filtered through a 40 μm cell strainer. Finally, live cells were passed through a Percoll gradient (35%/70%), recovered from the interface, washed and counted.

### Single cell barcoding and library preparation

Lamina propria cells were prepared for cell sorting as described above, using only LIVE/DEAD dye and an anti-CD45 (BioLegend, clone: 30-F11) fluorochrome-conjugated antibody.

Recovered CD45 positive cells were prepared for scRNAseq analysis using a 10X Genomics Chromium Controller. Single cells were processed with GemCode Single Cell Platform using GemCode Gel Beads, Chip and Library Kits (v2) following the manufacturer’s protocol. Libraries were sequenced on HiSeq 3000 (Illumina).

### Skin wounding and cell isolation

#### Skin wounding and tdRFP monocyte adoptive transfer

Monocytes were isolated from B6.RFP mice via Immuno-magnetic depletion of whole bone marrow by incubating samples with biotinylated antibodies against: CD3 (eBioscience, clone: 145-2C11), CD4 (eBioscience, clone: GK1.5), CD8 (eBioscience, clone: 53-6.7), CD45R (Biolegend, clone: RA3-6B2), CD19 (eBioscience, clone: eBio1D3), NK1.1 (eBioscience, clone: PK136), Ter119 (Biolegend), CD49b (Biolegend, clone: DX5), Ly6G (Biolegend, clone: 1A8) and CD117 (Biolgend, clone: 2B8). Anti-Biotin microbeads (Miltenyi) were then used according to the manufacturer’s protocol. Enrichment was validated by staining purified bone marrow monocytes with antibodies against CD115 (Biolegend, clone: AFS98), Ly6C (BD Bioscience, clone: AL-21).

Wounding and preparation of wound tissue was performed as previously described (*62*). Briefly, mice were anesthetized by i.p. injection of Ketanest/Rompun (Park Davis, Bayer). Back skin was shaved and full-thickness excisional wounds were created using a standard biopsy puncher (Stiefel). Mice were housed individually during the entire time course of healing. Monocytes from female B6.RFP mice were isolated as described above, counted and then 3x10^6^ cells were adoptively transferred into each previously wounded C57/BL6 recipient via intravenous injection, either 2 or 12 days after injury. Wounds were excised 4 or 14 days after injury.

#### Cell isolation

Excised wound tissue was sectioned with a scalpel, placed in DMEM with 30 µg/mL Liberase TM Research Grade (Roche Applied Science) and incubated at 37°C for 90 min (shaking).

Digested wound tissue was mechanically disrupted for 5 min using the Medimachine System (BD Biosciences). Cells were passed through 70 µm and 40 µm cell strainer and washed with 1% bovine serum albumin and 2 mM EDTA in PBS. Isolated cells were stained for flow cytometry and cell sorting as described above, using the following fluorochrome-conjugated antibodies: CD11b (eBioscience, clone: M1/70), F4/80 (eBioscience, clone: BM8 or AbD Serotec, clone: CI:A3-1), MHC-II (eBioscience, clone: M5/114.15.2), Ly6C (BD, clone: AL-21), CD45 (eBioscience, clone: 30-F11) and CD301b (Biolegend, clone: URA-1). Dead cells were excluded by labelling them with 50 ng/mL DAPI (ThermoFisher scientific).

#### Single cell barcoding and library preparation

Single macrophages from wounded skin were index-sorted (BD FACS Aria II SORP) into 384 well plates containing 0.5 µl of nuclease free water with 0.2% Triton-X 100 and 4 U murine RNase Inhibitor (NEB), spun down and frozen at −80°C. Libraries were prepared following the Smart-seq2 workflow (*63*). Briefly, after thawing, 0.5 µl of a primer mix were added (5 mM dNTP (Invitrogen), 0.5 µM dT-primer (C6-aminolinker-AAGCAGTGGTATCAACGCAGAGTCGACTTTTTTTTTTTTTTTTTTTTTTTTTTTTTTVN), 1 U RNase Inhibitor (NEB)). RNA was denatured for 3 minutes at 72°C and the reverse transcription (RT) was performed at 42°C for 90 min after filling up to 10 µl with RT buffer mix for a final concentration of 1x superscript II buffer (Invitrogen), 1 M betaine, 5 mM DTT, 6 mM MgCl2, 1 µM TSO-primer (AAGCAGTGGTATCAACGCAGAGTACATrGrGrG), 9 U RNase Inhibitor and 90 U Superscript II. After synthesis, the reverse transcriptase was inactivated at 70°C for 15 min. The cDNA was amplified using Kapa HiFi HotStart Readymix (Peqlab) at a final 1x concentration and 0.1 µM UP-primer (AAGCAGTGGTATCAACGCAGAGT) under following cycling conditions: initial denaturation at 98°C for 3 min, 23 cycles [98°C 20 sec, 67°C 15 sec, 72°C 6 min] and final elongation at 72°C for 5 min. The amplified cDNA was purified using 1x volume of hydrophobic Sera-Mag SpeedBeads (GE Healthcare) resuspended in a buffer consisting of 10 mM Tris, 20 mM EDTA, 18.5 % (w/v) PEG 8000 and 2 M sodium chloride solution. The cDNA was eluted in 12 µl nuclease free water and the concentration of the samples was measured with a Tecan plate reader Infinite 200 pro in 384 well black flat bottom low volume plates (Corning) using AccuBlue Broad range chemistry (Biotium).

For library preparation up to 700 pg cDNA was desiccated and rehydrated in 1 µl Tagmentation mix (1x TruePrep Tagment Buffer L, 0.1 µl TruePrep Tagment Enzyme V50; from TruePrep DNA Library Prep Kit V2 for Illumina; Vazyme) and tagmented at 55°C for 5 min. Subsequently, Illumina indices were added during PCR (72°C 3 min, 98°C 30 sec, 13 cycles [98°C 10 sec, 63°C 20 sec, 72°C 1 min], 72°C 5 min) with 1x concentrated KAPA Hifi HotStart Ready Mix and 300 nM dual indexing primers. After PCR, libraries were quantified with AccuBlue Broad range chemistry, equimolarly pooled and purified twice with 1x volume Sera-Mag SpeedBeads. This was followed by Illumina 50 bp paired-end sequencing on a Novaseq6000 aiming at an average sequencing depth of 0.5 mio reads per cell.

### Single cell RNA sequencing analysis

#### Adipose tissue pre-processing

Samples were demultiplexed and aligned using Cell Ranger 2.2 (10X genomics) to genome build GRCm38 to obtain a raw read count matrix of barcodes corresponding to cells and features corresponding to detected genes. Read count matrices were processed, analyzed and visualized in R v. 4.0.0 (*64*) using Seurat v. 3 (*36*) with default parameters in all functions, unless specified. Poor quality cells, with low total unique molecular identifier (UMI) counts and high percent mitochondrial gene expression, were excluded. Filtered samples were normalized using a regularized negative binomial regression (SCTransform) (*65*) and integrated with the reciprocal principal component analysis (rpca) approach followed by mutual nearest neighbors, using 50 principal components. Integrated gene expression matrices were visualized with a Uniform Manifold Approximation and Projection (UMAP) (*66*) as a dimensionality reduction approach.

Resolution for cell clustering was determined by evaluating hierarchical clustering trees at a range of resolutions (0 - 1.2) with Clustree (*67*), selecting a value inducing minimal cluster instability. Datasets were subsetted to include only macrophages, based on the expression of key macrophage markers (*Adgre1*, *Csf1r*, *H2-Ab1*, *Cd68*, *Lyz2*, *Itgam*, *Mertk*), retaining only 500 randomly selected cells per biological condition. Macrophage only datasets were then split along conditions, and processed anew as described above, to obtain a reference macrophage dataset.

#### Differential gene expression, pathway enrichment analysis and gene set score calculation

Differentially expressed genes between clusters were identified as those expressed in at least 40% of cells with a greater than +1 log fold change and an adjusted p value of less than 0.01, using the FindMarkers function in Seurat v.3 with all other parameters set to default. Ribosomal protein genes were excluded from results.

Cluster specific genes were explored for pathway enrichment using StringDB (*68*), where characteristic gene sets were mapped to specific functions (Supplemental Table 2). Gene set scores were calculated using the AddModuleScore function in Seurat v.3 with default parameters. Briefly, the average expression levels of each identified gene set was calculated on a single cell level and subtracted by the aggregated expression of randomly selected control gene sets. For this purpose, target genes are binned based on averaged expression, and corresponding control genes are randomly selected from each bin.

#### Trajectory inference, pseudotime calculation and trajectory dependent gene regulation

Macrophage activation trajectories and pseudotime estimations were calculated with slingshot v. 1.6.1 (*30*), using UMAP projection and pre-calculated clustering as input for getLineages and getCurves functions with default parameters, setting origin to cluster 4. Cells in the reference dataset were then assigned an activation trajectory and corresponding pseudotime value.

Trajectory dependent gene regulation was calculated first by extracting the 2000 genes with the most variance in expression in cells participating in each detected pathway, then fitting general additive models (GAM) to these genes and extracting the coefficient’s p value using gam v. 1.20 (*69*). Fitted models had the expression of each gene as a response variable (G) and pseudotime (t) as an independent variable using locally estimated scatterplot smoothing (loess) smooth terms (lo):

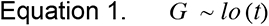

Extracted coefficient p values were adjusted for multiple comparisons using a false discovery rate correction with the p.adjust function from stats v. 4.0.0 (*64*). Adjusted p values were used to rank genes, and an arbitrary threshold was used to select most significant model fits.

#### Query dataset retrieval and preprocessing

Lamina propria samples containing CD45^+^ cells were demultiplexed, mapped, filtered, clustered and projected as described above, maintaining individual biological conditions separate. A “Macrophage score” (Supplemental Table 2) was calculated for this dataset as described above, and clusters containing macrophages extracted.

Remaining query datasets available from public repositories were retrieved (Supplemental Table 3), obtaining matrices with raw unfiltered read counts for detected genes in barcoded cells. Low quality cells were removed and then datasets were subsequently clustered and projected as specified above. Where non-macrophage cells were included in the sequencing experiment, a “Macrophage score” (Supplemental Table 2) was calculated, and these cells extracted, keeping at most 500 randomly selected cells per biological condition.

#### Data imputation and label transfer

Query datasets were individually normalized with SCTransform, then integrated within tissues using the rpca approach as described above. Each resulting integrated dataset was then compared to the reference dataset to transfer identified labels and harmonize data via imputation using the FindTransferAnchors (dims = 50, npcs = 50, k.filter = 5, max.features = 100, k.anchor = 5) and TransferData (dims = 30, k.weight = 25, sd.weight = 1) functions from Seurat v.3 with the specified parameters. These were benchmarked using two negative control datasets containing multiple types of immune cells, selecting values reducing the number of low quality anchors and increasing the label probability score. A threshold of 80% (0.8) was set for the label probability based on negative controls to consider label allocation as successful.

Additionally, label probability distributions across clusters in query datasets were investigated to evaluate goodness of transfer. Where a cluster was found to be dominated by a single label, and a portion of the cells within the cluster had a high label probability score, the corresponding label was assigned to the entire cluster.

#### Differential gene expression analysis within biological conditions

Differentially expressed genes between biological conditions within a particular macrophage activation stage were identified as those expressed in at least 25% of cells with a greater than +0.25 log fold change and an adjusted p value of less than 0.01, using the FindMarkers function in Seurat v.3 with all other parameters set to default. Gene expression differences were based on normalized data, not on imputed data. Ribosomal protein genes were excluded from results.

#### Wounded skin pre-processing, label transfer and analysis

Raw reads were mapped to the mouse genome (GRCm38) and splice-site information from Ensembl release 87 with gsnap v.2018-07-04 (*70*). Uniquely mapped reads and gene annotations from Ensembl release 87 were used as input for featureCounts v. 1.6.2 (*71*) to create counts per gene and cell. Filtering, clustering and projection was performed as described above, with the addition of filtering based on reads mapping to ERCC spike-in controls. Label and data transferred was performed as described above.

Flow cytometry data associated with individually barcoded cells was used to calculate a UMAP projection using uwot (*72*), which was then clustered with kmeans v. 4.0.0 (*64*), setting k based on the total within cluster sum of squares.

#### Cross-condition data integration and marker selection

Cells in each dataset with a high label probability score were extracted, split by tissue retaining only the depth corrected RNA counts and label assignment, then normalized using SCTransform. These datasets were then integrated, clustered and projected as described above, without giving priority to the reference data, which was further subsampled (500 randomly selected cells) prior to integration. Differentially expressed genes between macrophage activation stages were identified as those expressed in at least 25% of cells with a greater than +0.25 log fold change and an adjusted p value of less than 0.01, using the FindMarkers function in Seurat v.3 with all other parameters set to default. Surface expression of genes was determined based on GO annotation (Cell surface – GO:0009986).

#### Transcriptional network, transcription factor and pathway enrichment analysis

Genes, for which a significant association between pseudotime and gene expression was found, were analysed using StringDB (*68*) to build a network of protein-protein interactions based on experimental evidence, reported co-expression and database mining. Resulting undirected network was retrieved and analysed using igraph v. 1.2.5 (*73*) and ggraph v. 2.0.2 (*74*). Weight of edges connecting nodes was calculated as follows:

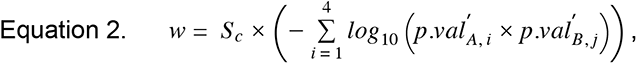

where *w* is the edge weight, *S_c_* is the confidence score, *p.val* is the scaled adjusted p value for the pair of connected genes A and B in activation path *i*, with *i* ranging through the four defined activation paths. Adjusted p values were scaled within each activation path to center the values around 0 making them more comparable across paths. Low weight edges and disconnected nodes were filtered, and resulting network nodes were annotated for betweenes, degree, eigen-centrality and strength using native function provided in igraph. The Fruchterman-Reingold algorithm was used to calculate the network’s layout. Network clustering was performed using the cluster_louvain function (*75*).

Network clusters were interrogated for pathway enrichment using either Biological process Gene Ontology annotation or KEGG pathways with goseq v. 1.40.0 (*76*). Gene set scores (Supplemental Table 2) for top enriched pathways were calculated as specified above and the pathway score variance calculated across cells within each activation trajectory. Resulting variances were used for heatmaps as a proxy for enriched pathway regulation within each activation trajectory.

Transcription factor enrichment analysis was performed with RcisTarget v. 1.8.0 (*77, 78*). The number of incidences of each identified transcription factor across all clusters was counted, and visualized as a word cloud using wordcloud2 v 0.2.1 (*79*).

### Quantification and statistical analysis

Statistical analysis was performed in R v. 4.0.0 (64), using functions from the base stats package to calculate a single factor anova with aov followed by Tukey Honest Significant Differences to determine statistically significant differences between means (Figure 2G-H) or only a single factor anova to establish significant clustering effects (Figure 5M). Ex vivo results are represented as dots for individual mice. Selection of sample size was based on extensive experience with similar assays.

## Acknowledgements

The authors would like to thank Dr. Paul Gueguen, Dr. Beth Kelly, members of the Pearce laboratories, the staff at the DeepSequencing and Bioinformatics facilities at the MPI-IE, as well as the members of the single cell unit from the DRESDEN-concept Genome Center for technical support and constructive criticism. This study was supported by the NIH (AI110481 to E.J.P.), the Max Planck Society and the German Research Foundation (Leibniz Prize to E.L.P.; Germany’s Excellence Strategy: CIBSS EXC-2189 Project ID 390939984 to E.L.P. and CECAD EXC-2030 project ID 390661388 to S.A.E.; Research Unit FOR2599 to S.A.E., A.D., A.R., P.J.M. and E.J.P.; Collaborative Research centres: CRC1218 project ID 269925409 and CRC1403 project ID 1403-414786233 to S.A.E.). A.M.K., A.C. and S.D. were supported by the Alexander von Humboldt Fellowship Foundation.

## Author contributions

Conceptualization, D.E.S., S.A.E., A.G., A.R., P.J.M. and E.J.P.; Methodology, D.E.S.; Software, D.E.S.; Formal analysis, D.E.S.; Investigation, D.E.S., Y.G., E.M., A.M.K., K.M.G., A.C., G.C., A.G., A.D., S.W., S.R., and J.D.C.; Data Curation, D.E.S. and Y.G.; Writing – Original Draft, D.E.S. and E.J.P.; Writing – Review & Editing, D.E.S., S.W., S.D., E.L.P., S.A.E., A.G., A.R., P.J.M. and E.J.P.; Funding Acquisition, E.L.P., S.A.E., A.G., A.R., P.J.M. and E.J.P.; Visualization, D.E.S.; Supervision, E.J.P.

## Supplemental Materials

**Figure S1.**
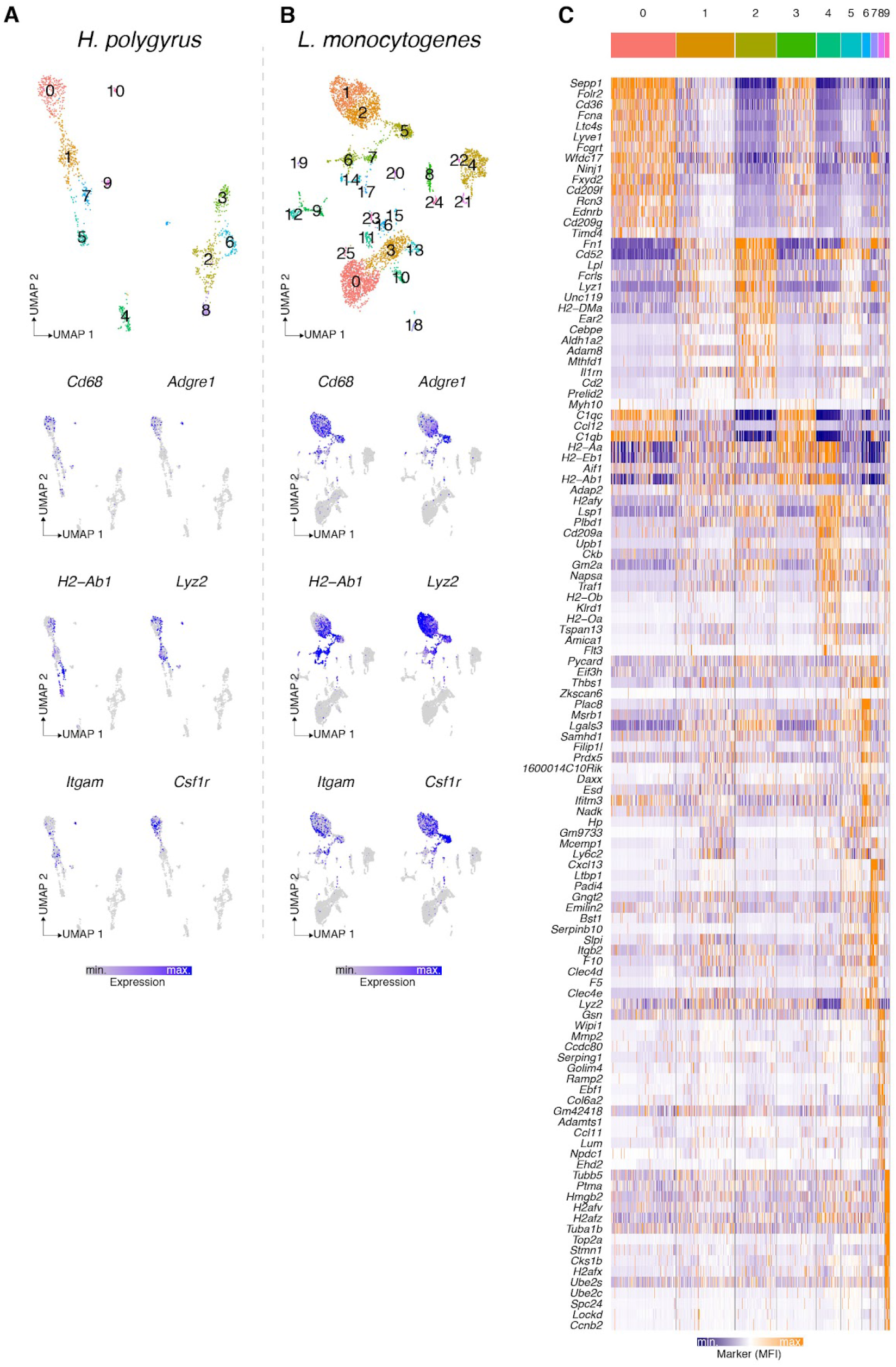
Related to Figure 1. (A) UMAP of CD45^+^ SVF cells isolated from mesenteric adipose tissue of naïve and *H. poly* infected mice, labelled and colorer according to identified clusters (top) or showing relative expression (low - gray; high - blue) of macrophage marker genes (bottom). (B) UMAP of SVF cells isolated from popliteal adipose tissue of naïve and *L. mono* infected mice, labelled and colorer according to identified clusters (top) or showing relative expression (low - gray; high - blue) of macrophage marker genes (bottom). (C) Heatmap showing relative expression (low - blue; high - orange) of top significantly (adjusted p value < 0.01) regulated genes (log fold change > 0.25) genes across identified clusters in Figure 1B.

**Figure S2.**
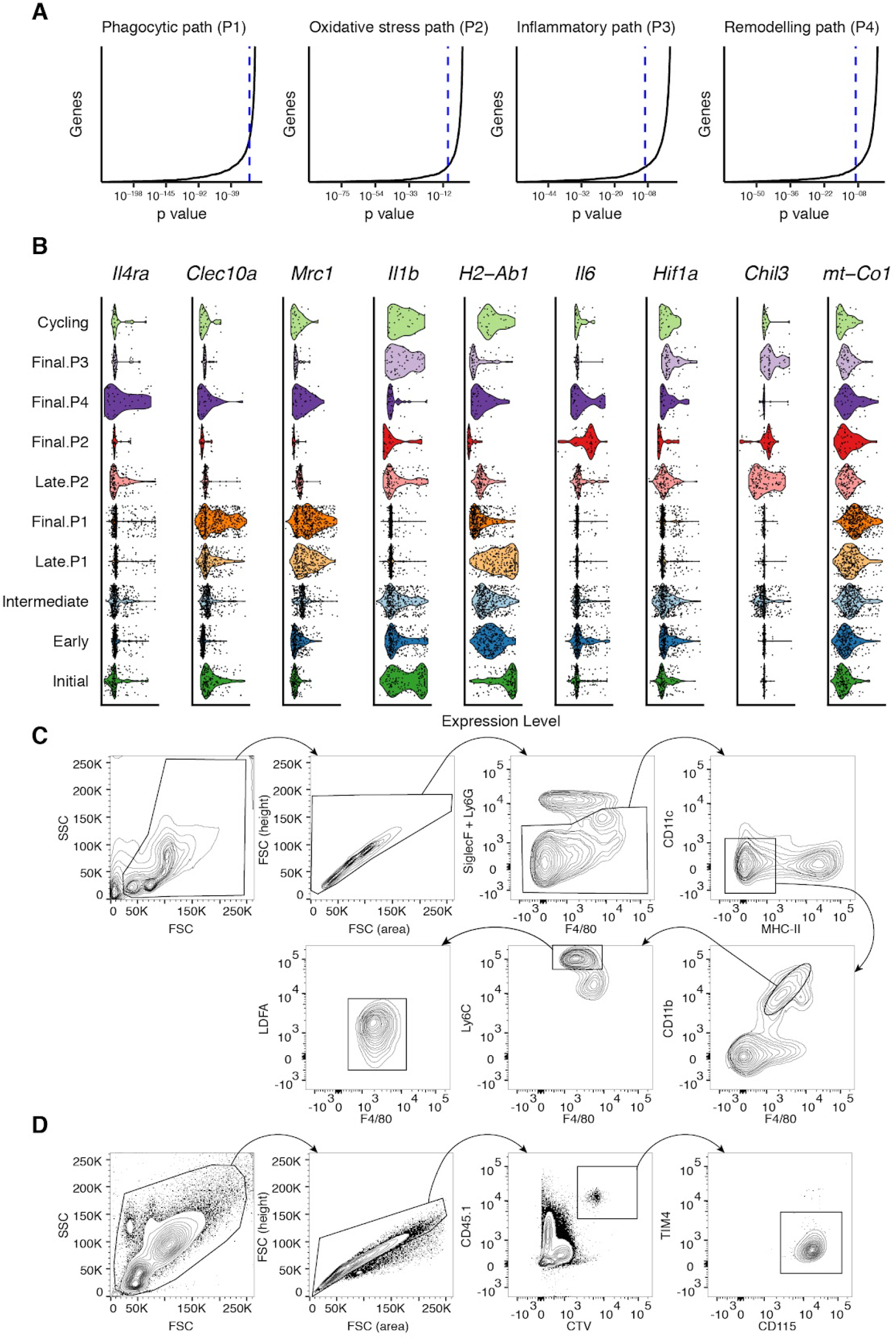
Related to Figure 2. (A) Adjusted p value distribution for GAM fits for each activation path, showing an arbitrary significance threshold (blue dashed line - 1x10^-9^). (B) Violin plots showing relative gene expression of macrophage activation markers across identified stages. (C-D) Representative flow cytometry plots showing gating strategy for the isolation of live bone marrow monocyte precursors (C), or analysis of adoptively transferred cells in the peritoneal cavity (D).

**Figure S3.**
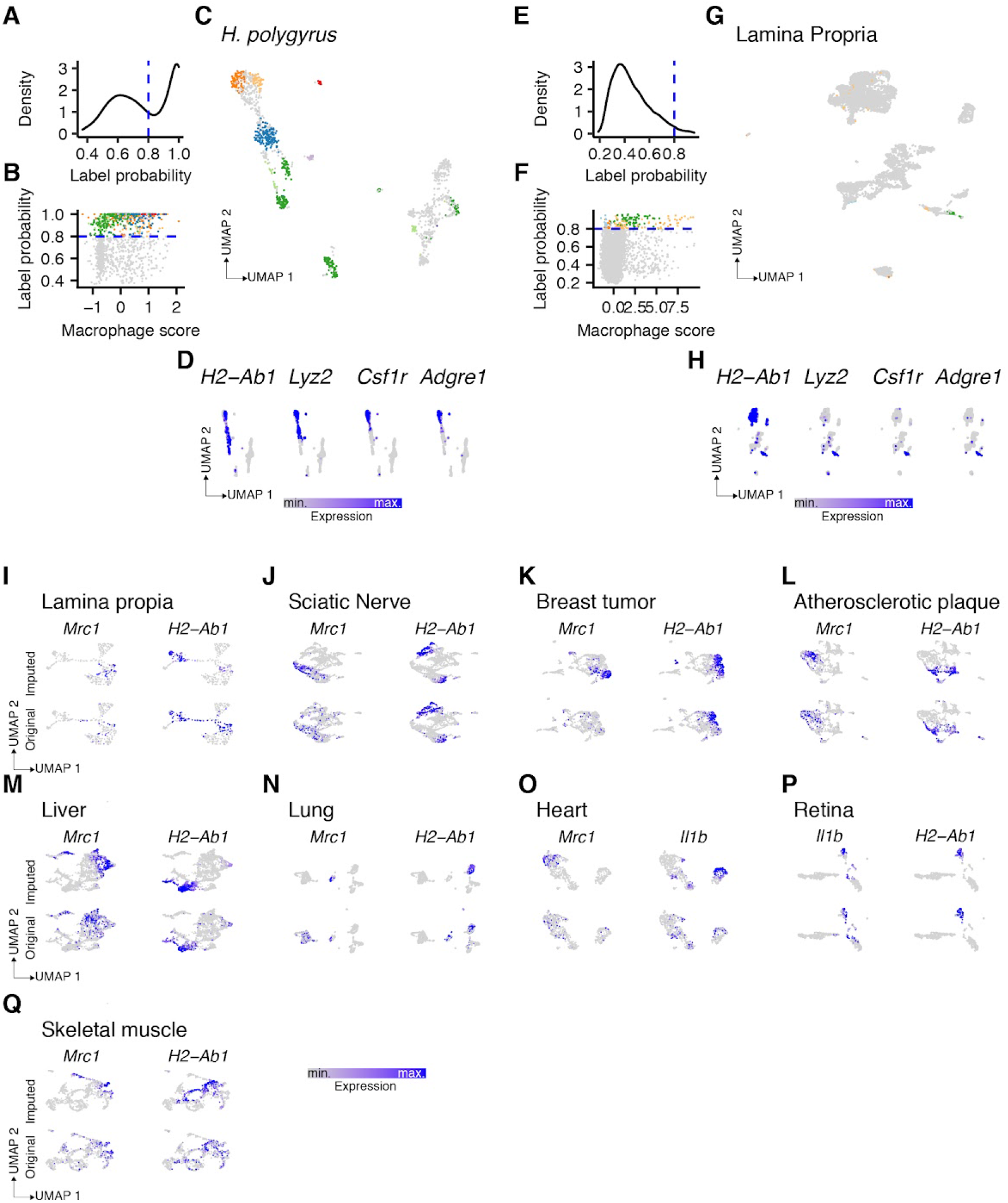
Related to Figure 3. (A-H) Analysis of CD45^+^ cells from either the SVF of *H. poly* infected animals (A-D) or the lamina propria (E-H). (A, E) Label probability distribution showing confidence threshold (dashed blue line) for label assignment. (B, F) Relationship between label probability and macrophage score in cells shown as dots and colored according to transferred labels. Dashed blue line indicates confidence threshold (80%) for label transfer. (C, G) UMAP showing assigned labels (colored cells) and cells with no classification in grey. (D, H) UMAP showing relative expression (low - gray; high - blue) of macrophage specific genes. (I-Q) UMAP showing relative expression (low - gray; high - blue) of 2 genes per indicated dataset, illustrating the effects of data imputation procedure by comparing expression patterns in original (bottom) and imputed (top) data.

**Figure S4.**
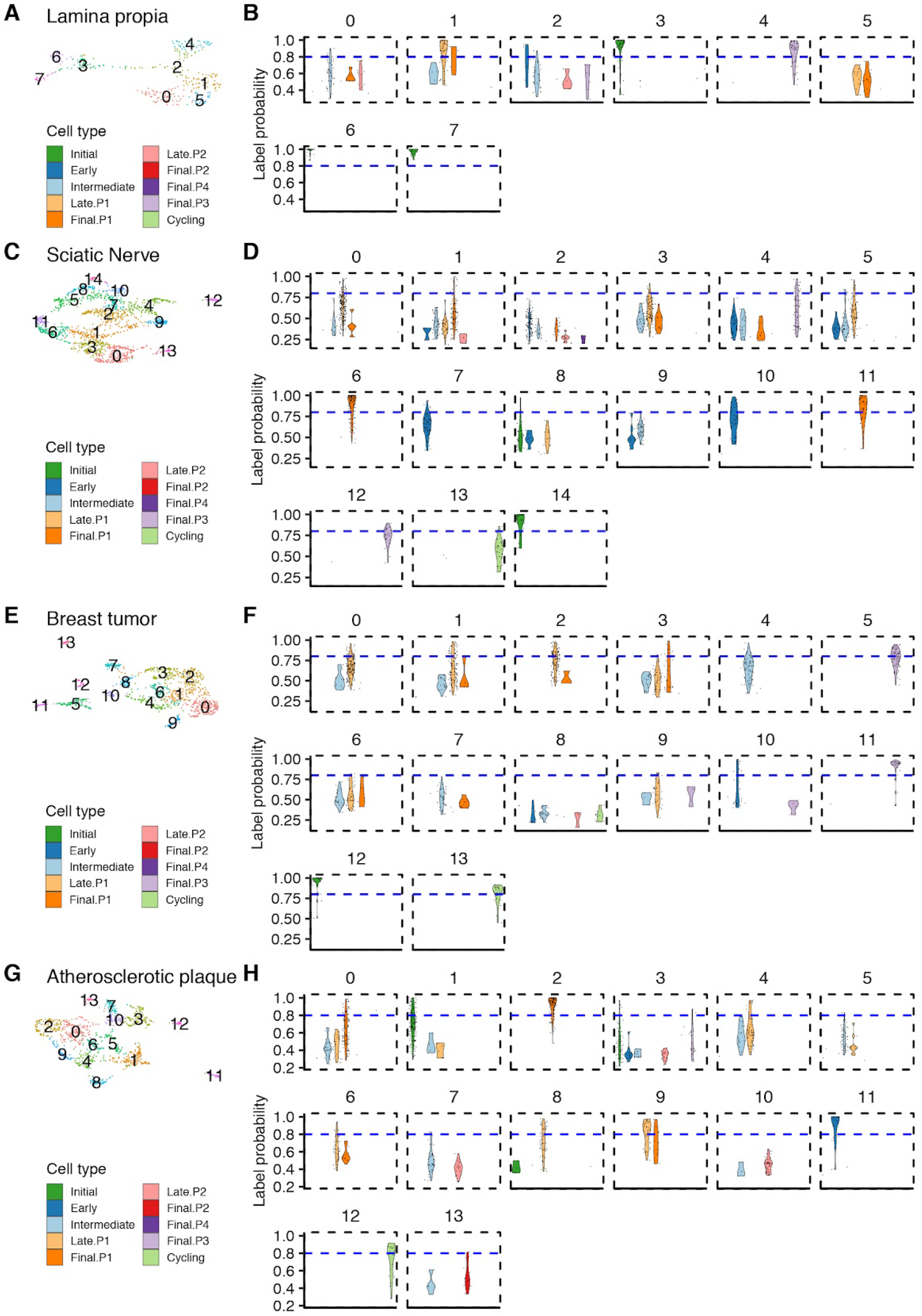
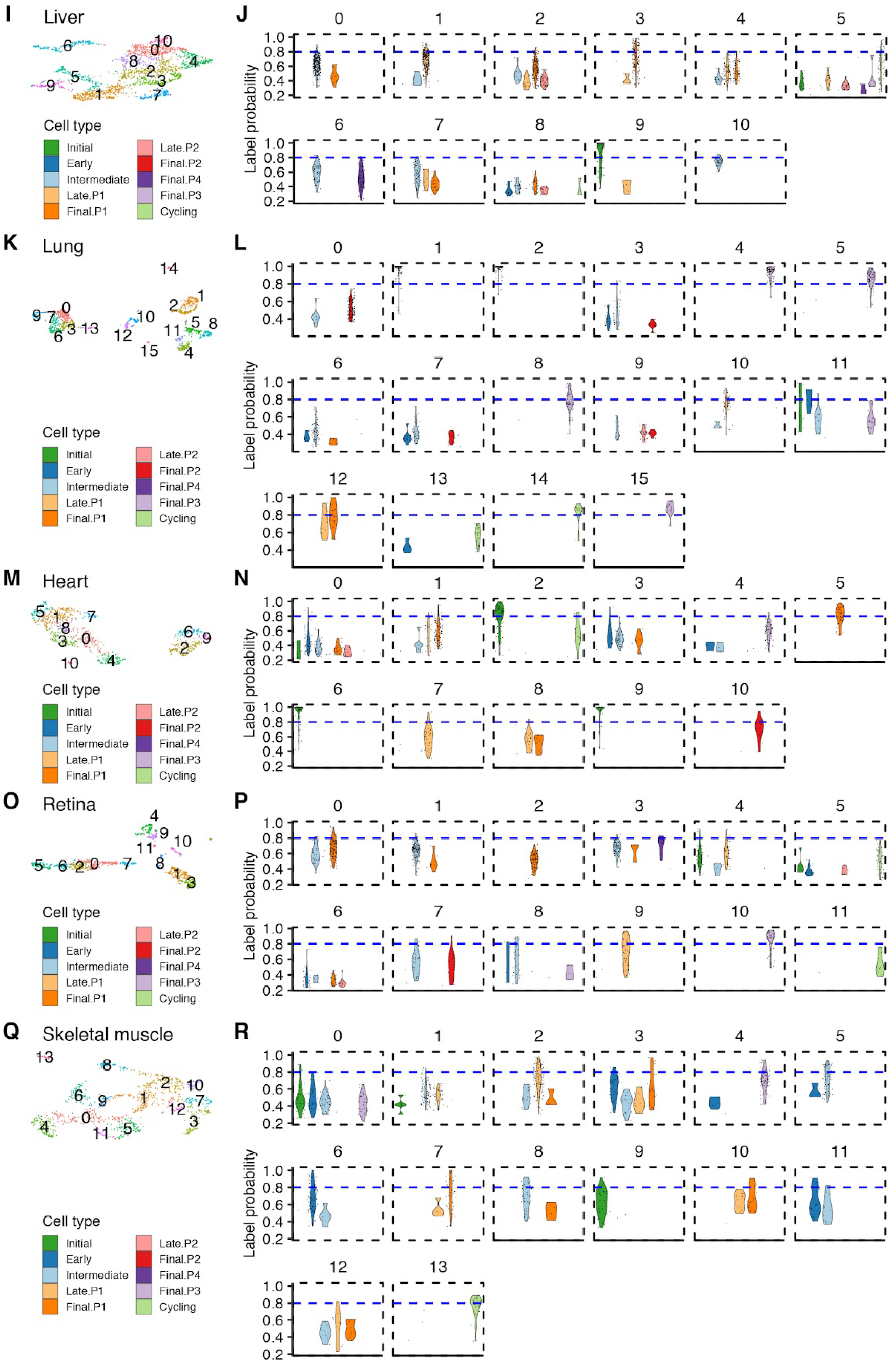
Related to Figure 3. (A-R) Query datasets were clustered and visualized as UMAPs (A, C, E, G, I, K, M, O, Q) and violin plots illustrating the label probability distribution for every detected label within each identified cluster (B, D, F, H, J, L, N, P, R). Dashed blue line indicates confidence threshold (80%) for label transfer.

**Figure S5.**
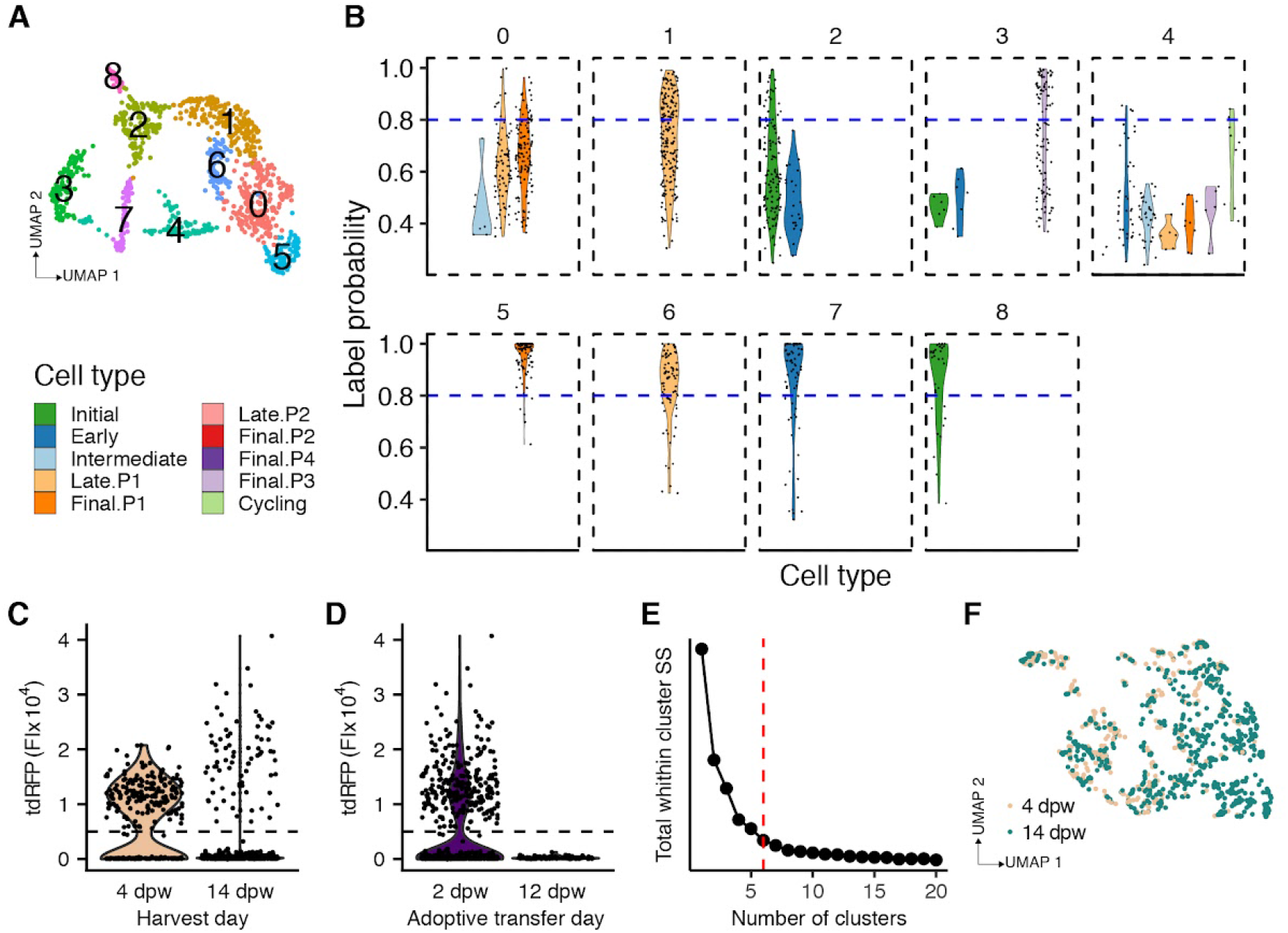
Related to Figure 5. (A) UMAP of wounded skin macrophages indicating identified clusters. (B) Violin plots illustrating the label probability distribution for every detected label within each identified cluster. Dashed blue line indicates confidence threshold (80%) for label transfer. (C-D) tdRFP+ fluorescence intensity (FI) in all cells across harvest day (C) or adoptive transfer day (D). (E) Total within cluster sum of squares (SS) as a function of k means (number of clusters). Dashed red line indicates the selected number of clusters (k= 6). (F) UMAP calculated based on indexed flow cytometry data with day of tissue harvest indicated.

**Figure S6.**
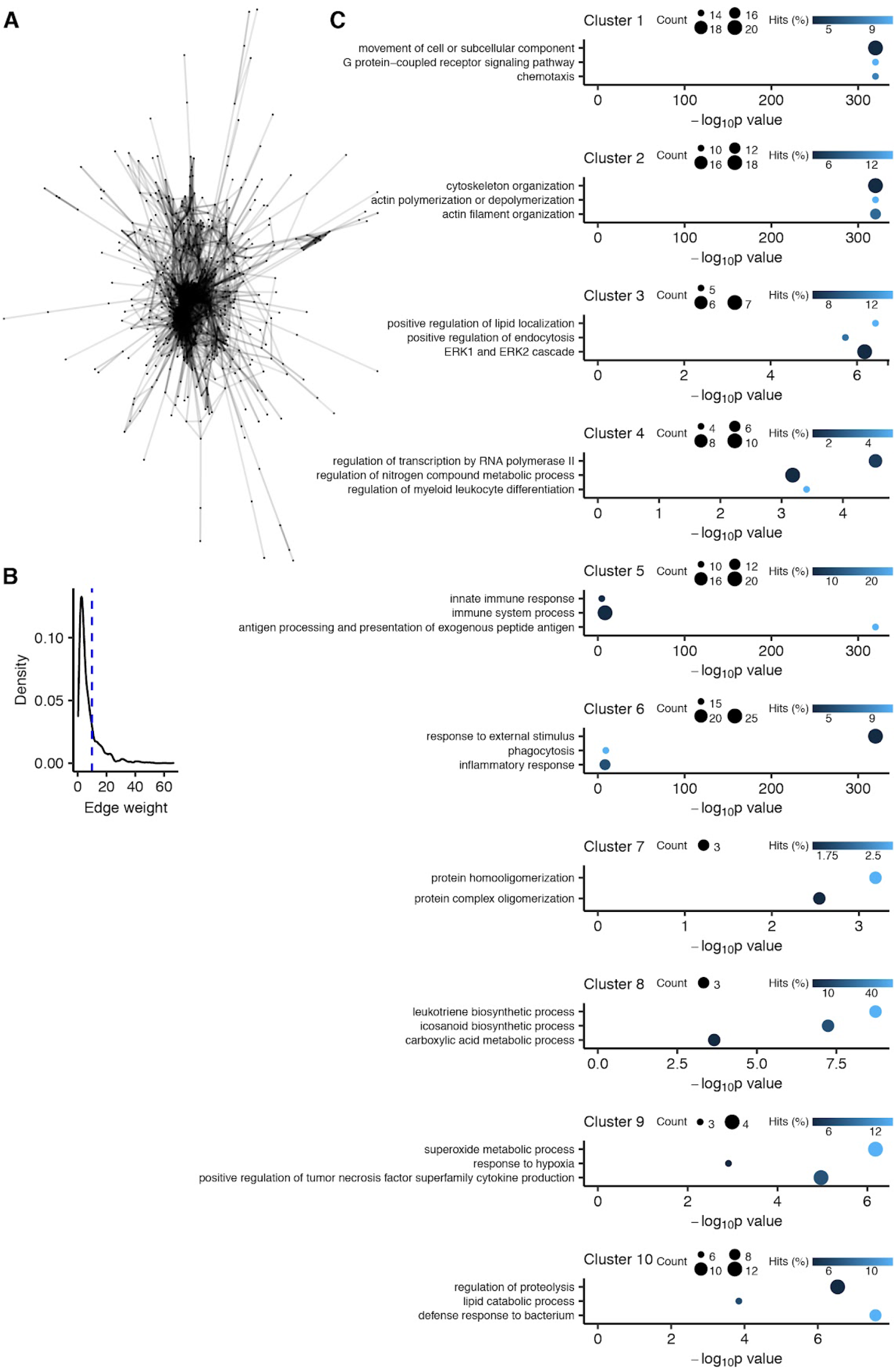
Related to Figure 7. (A) Transcriptional network or protein-protein interactions, depicting genes as nodes (n = 586) and interactions as edges (n = 3355). (B) Edge weight distribution indicating threshold (dashed blue line) for edge trimming. (C) Dot plot depicting top enriched GO terms for each identified network cluster.

**Supplemental Table 2.**
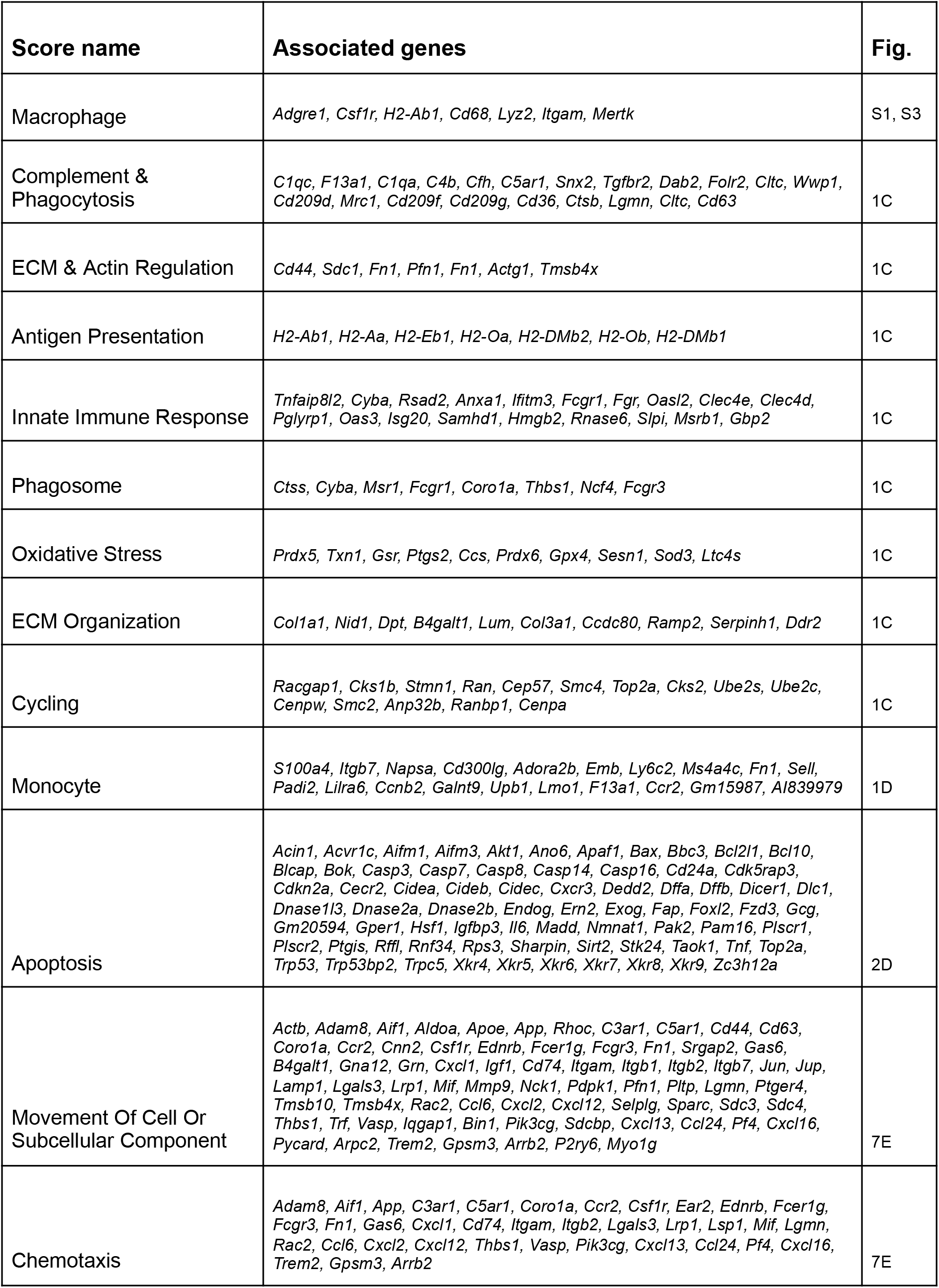
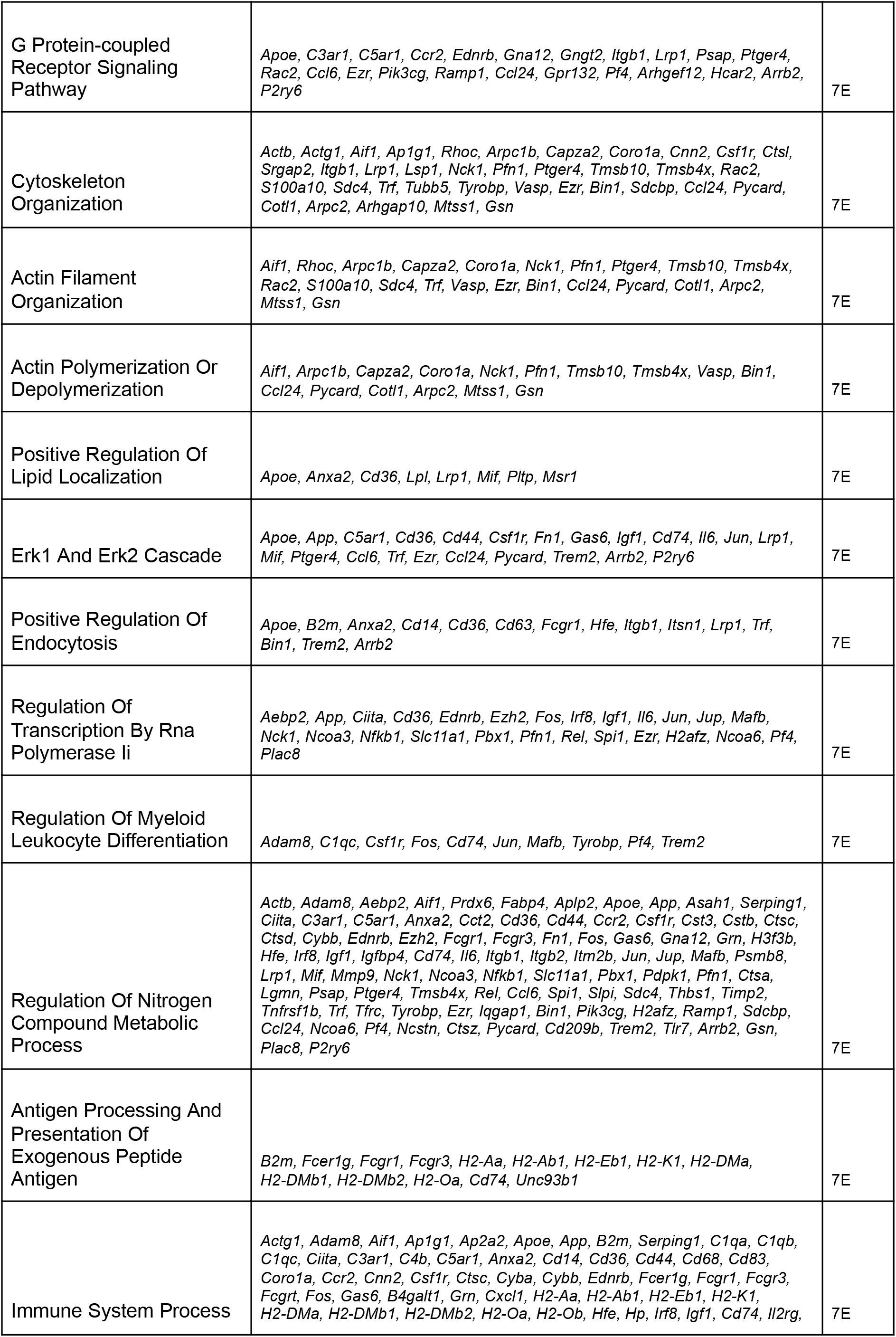
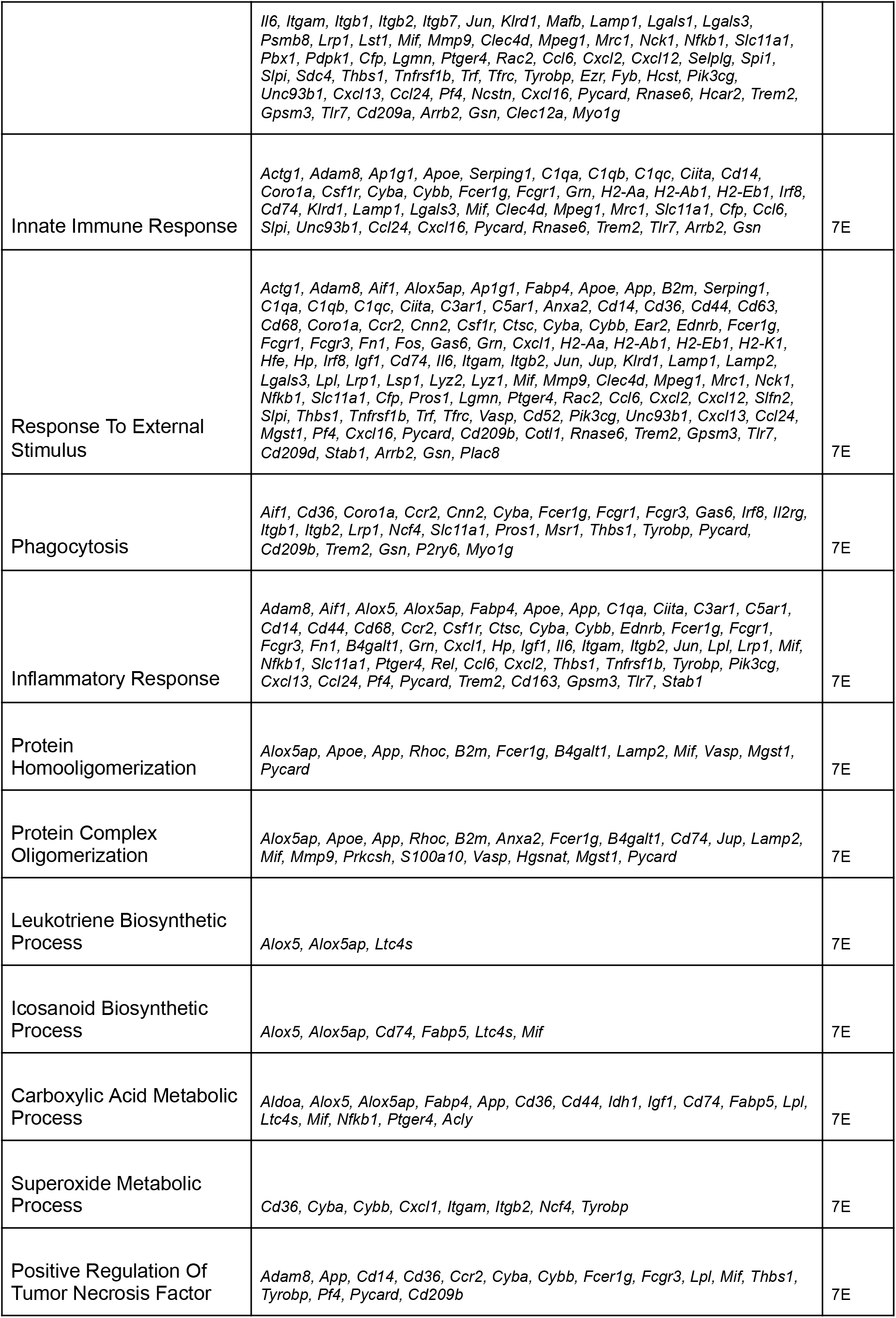
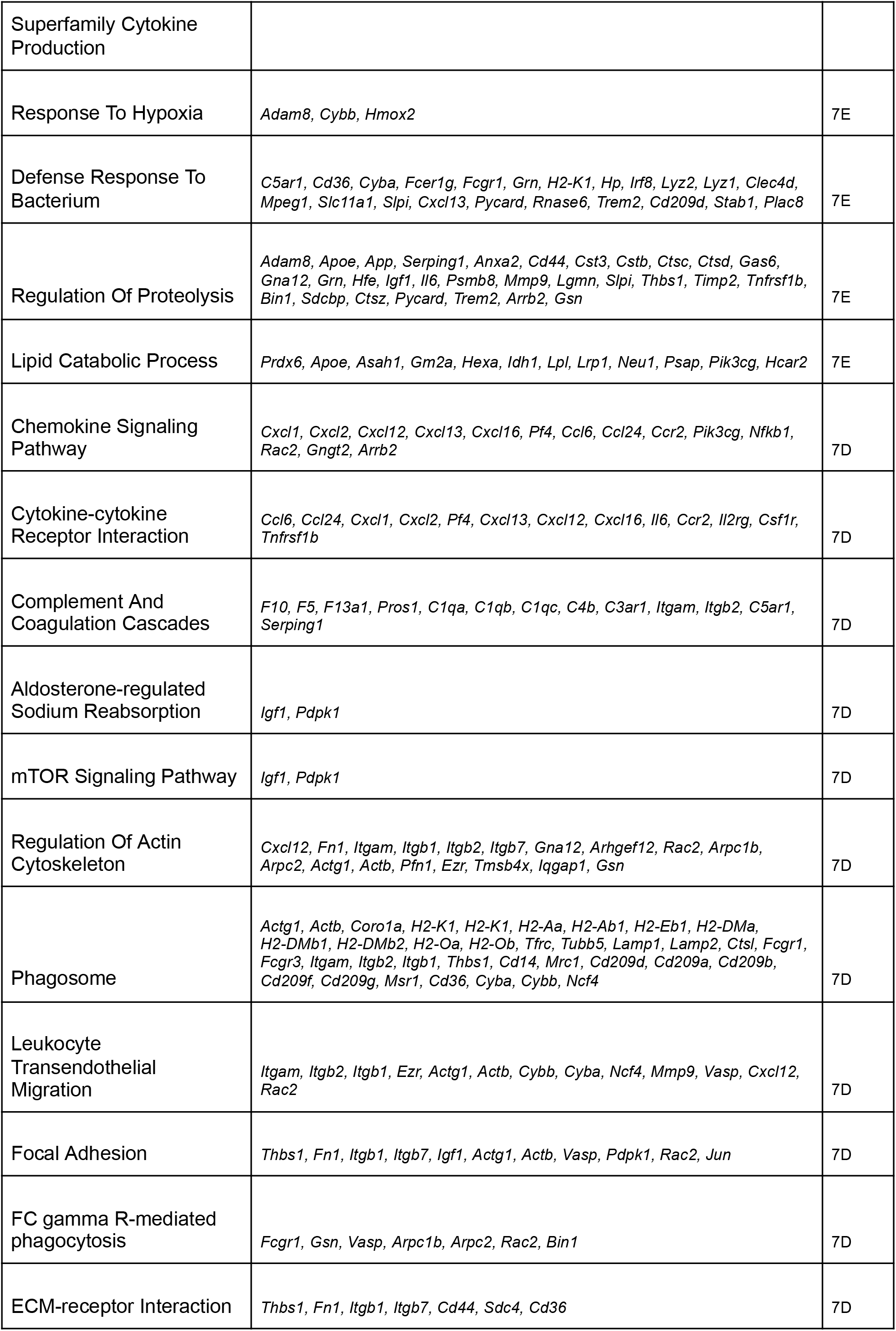
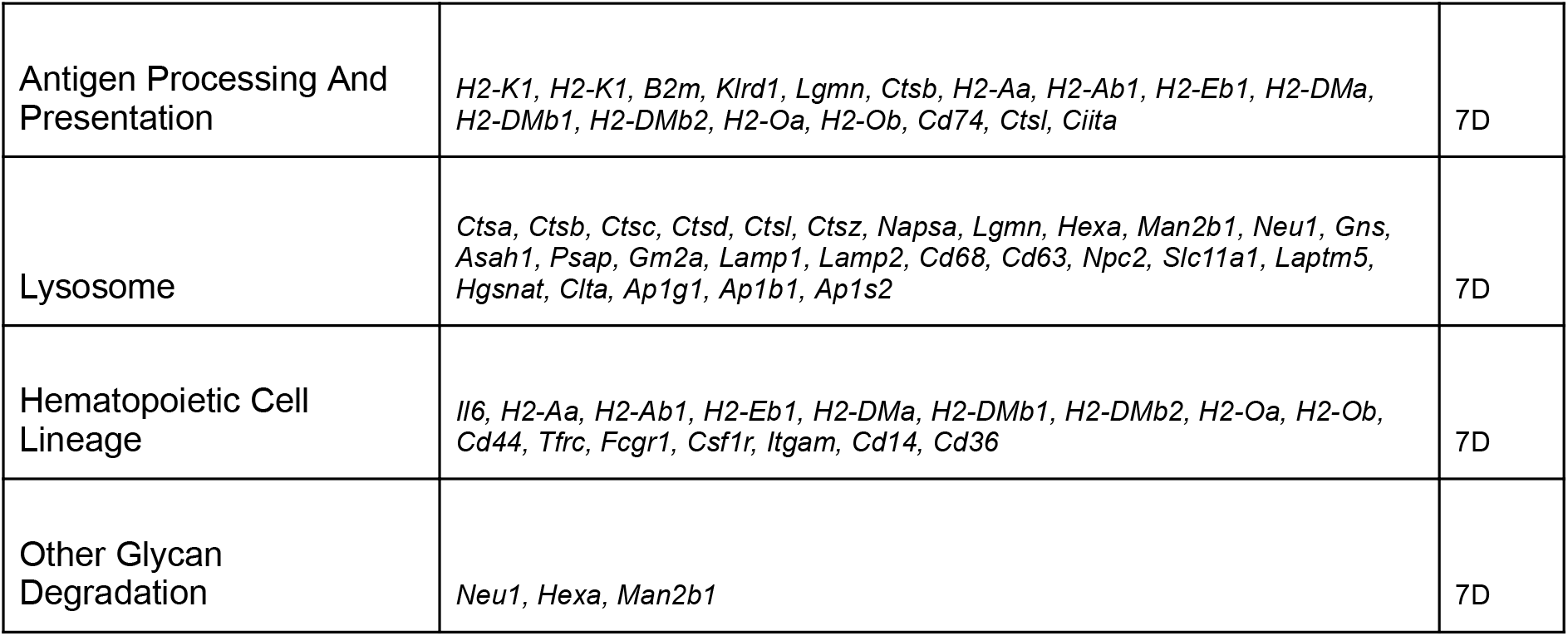
Gene set scores. Genes used in calculating all gene set scores shown in the study.

**Supplemental Table 3.** Dataset description and source. A description of single cell RNA-seq datasets used in the study, including a brief outline of the immunological conditions within each dataset, is presented. Accession codes and associated publications are also given.

| Tissue | Cells | Conditions | Description | Timing | Source | Accession | Technology |
| --- | --- | --- | --- | --- | --- | --- | --- |
| Popliteal adipose | Stromal vascular fraction | Naïve control or *L. monocytogenes* infection | Footpad s.c. injection of *L. monocytogenes* | 24 hpi | This publication | GSE171328 | 10X Chromium controller |
| Mesenteric adipose | Stromal vascular fraction | Naïve control or *H. polygyrus* infection | Oral infection with *H. polygyrus* larvae | 14 dpi | This publication | GSE157313 | 10X Chromium controller |
| Lamina propria | CD45+ cells | Control (chow) diet or High fat diet | Rodent Diet 60% kcal from fat | 12 wks | This publication | GSE171330 | 10X Chromium controller |
| Sciatic nerve | Macrophages | Naïve control or Nerve crush | Sciatic nerve was surgically exposed then crushed. | 1 and 5 dpw | Ydens et al. 2020 | GSE144707 | 10X Chromium controller |
| Breast tumor | Myeloid cells | Spontaneous tumor in mice with WT or *Dab2* deficient macrophages | Dissected spontaneous lobular breast carcinoma in MMTV-PyMT *Dab2*<sup>fl/fl</sup> Tie2-cre+ or Tie2-cre- mice | 13 wks | Marigo et al. 2020 | GSE152674 | 10X Chromium controller |
| Atherosclerotic plaque | Macrophages | Progressing or Regressing lesion | 20 weeks western diet or 18 weeks western diet then 2 weeks chow diet with apolipoprotein B antisense oligonucleotide treatment (50 mg/kg, 2 doses/wk) | 20 wks | Lin et al. 2019 | GSE123587 | 10X Chromium controller |
| Lung | CD45+ cells | Naïve control or *Cryptococcus neoformans* infection | Oro-tracheal exposure to *Cryptococcus neoformans* yeasts | 9 hpi | Xu-Vanpala et al. 2020 | GSE146233 | 10X Chromium controller |
| Liver | Non-parenchymal liver cells | Healthy control or Fibrotic tissue | Overnight fasting followed by o.g. administration of carbon tetrachloride in corn oil (1:4) or vehicle control twice per week | 2 and 4 wks | Terkelsen et al. 2020 | GSE145086 | 10X Chromium controller |
| Heart | CD45+ cells | Healthy control or Infarcted tissue | Left anterior descending coronary artery occlusion by permanent suture ligation. | 4 days | King et al. 2017 | GSE106472 | inDrop (Zilionis et al, Nature Protocols 2016) |
| Retina | CD45+ cells | Healthy (dark) control or Neurodegeneration (light) | Arrestin 1 deficient maintained in constant darkness before exposure to light (200 lux, 48 hours). | 48 hours | Ronning et. al. 2019 | GSE121081 | 10X Chromium controller |
| Skeletal muscle | CD45+ cells | Naïve control or *Toxoplasma gondii* infection | Oral infection with ME49 *T.gondii* cysts | 28 dpi | Jin et. al. 2018 | GSE113111 | 10X Chromium controller |
| Brain | Microglia | Steady state | No intervention | E14, P4/5, P30, P100 | Hammond et. al. 2019 | GSE121654 | 10X Chromium controller |
| Skin | Macrophages | Wounded skin | Sterile wounding via skin punch biopsies. i.v. administration of tdRFP+ monocytes. | 4 or 14 dpw | This publication | GSE | Smart-Seq2 |

